# Animals or plants? Evolutionary branching of sessile versus mobile cognitive agents in noisy environments

**DOI:** 10.64898/2026.09.08.750148

**Authors:** Jordi Pla-Mauri, Ricard Solé

## Abstract

The evolution of complex cognition has been tied to movement and the need to locate resources in uncertain environments. Plants present a striking contrast: despite sophisticated environmental responses, their sessile lifestyle involves predictable energy capture and growth-mediated foraging. What ecological conditions could separate these alternative strategies? We address this question using spatially explicit individual-based simulations and Adaptive Dynamics models in which evolving agents allocate resources between harvesting a stable energy source (light) and exploiting patchy, fluctuating resources through costly sensing and movement. Starting from intermediate generalists, evolution repeatedly produces two contrasting specialist lineages: sessile, light-dependent agents that abandon sensing and locomotion, and mobile foragers that sacrifice light harvesting while investing in active resource search. Spatial gradients promote divergence through niche partitioning, but specialization also emerges in homogeneous environments through frequency-dependent ecological feedbacks, showing that environmental heterogeneity is not required. Under linear or convex returns, intermediate strategies can be disadvantaged because they bear the costs of both resource-acquisition systems without fully exploiting either; under sufficiently concave returns, intermediate strategies may instead be favored. These results suggest that plant- and animal-based organizations can emerge as alternative evolutionary solutions to a common foraging problem. More broadly, they support the idea that the evolution of costly cognitive machinery depends critically on the statistical structure of the resources that organisms must find and exploit.

## I. INTRODUCTION

The explosive rise of multicellular diversity at the Cambrian boundary marked the development of a biosphere where organismal complexity moved beyond simple, passive behavioral patterns to embodied complex cognitive agents [1, 2]. Once in place, neural systems and the potential for movement allowed cognitive complexity to flourish [3–5]. Arms races, where other species became an intrinsic part of evolutionary dynamics [6, 7], boosted different strategies to find available resources and adapt to their fluctuations over time. That is a standard picture for metazoans. As soon as bilaterians, with a well-defined bilateral symmetry, made their first appearance, nervous systems soon followed, eventually evolving a distinct central control system and becoming prediction machines [8].

Two crucial innovations were movement [9] and sight [10, 11]. Biologically, the appearance of complex eyes did not just add a new sense; it restructured the entire evolutionary game. Predation, camouflage, signaling, speed, spatial strategy, and arms races exploded at once. Information about the world suddenly traveled at light speed across distance. The so-called *moving hypothesis* [12] posits that such evolved exploration of habitats and their necessary sensors paved the way for brains. An important consequence of this idea is that prediction was both a cause and a consequence of animal movement. While perception and movement implied some metabolic constraints [13, 14], the reward of finding rich resources in a fluctuating environment can largely counterbalance the cost of a complex cognitive apparatus.

But what about plants? Are they also equipped with a similar cognitive potential despite their sessile nature? As autotrophs, plants have evolved a complex molecular machinery that gathers light and transforms it into energy and matter. This energy source does not require movement beyond the morphological changes associated with shading responses and seed dispersal. Does such a constraint limit the potential cognitive repertoire of plants? Can *plant intelligence* [15–17] compare with the capacities exhibited by animals? Plants certainly gather information from their environment, reacting to its changes by means of multiple plastic responses [18– 20]. But this information is integrated to control physiological and developmental processes [21] to a large extent toward growth and defense [22, 23]. Despite the use of neural-like analogies, from those in the domain of so-called *plant neurobiology* [24–26] to plant perceptrons [27], it is important to understand that analogies can easily fail [28]. Plants have no neurons or brains, and the perceptron analogy, grounded in a hardwired architecture, does not compare to the classical neural network [29].

It can be argued that plants are exceptionally well-suited to their “static” world by means of a completely different set of adaptive responses that do not require neurons nor cognition, and indeed, their survival strategies are based on traits that are rather orthogonal to the animal counterparts. Nevertheless, we can talk about *behavior* [30, 31] and *foraging behavior* in plants, which means that even non-moving agents can explore their environments in adaptive ways. This is clearly illustrated by how plant leaves and roots deal with multiple soil cues [32–34]. How does this compare with the foraging strategies of animals? Considerable controversy has been piling up over the last decade concerning this question [35].

An additional ingredient to consider is the rarity of intermediate forms combining heterotrophic and autotrophic functions. Carnivorous plants provide an obvious example [36–38]. Carnivory has evolved repeatedly in plants that inhabit nutrient-poor environments, where photosynthesis supplies carbon, but growth is limited by nitrogen, phosphorus, or other mineral nutrients. Capturing and digesting animal prey alleviates these limitations, but requires costly morphological and physiological adaptations and is advantageous only within a restricted range of ecological conditions. Carnivorous plants therefore occupy a narrow region of ecological and phenotypic space rather than constituting a general bridge between autotrophy and heterotrophy. At the opposite end of the spectrum are multicellular heterotrophs, such as “solar-powered” sacoglossan mollusks, that exploit photosynthesis through kleptoplasty [39–41]. These animals acquire chloroplasts from ingested algae and retain them within their tissues, where they can remain photosynthetically active for extended periods. However, kleptoplasty supplements heterotrophic metabolism rather than producing true autotrophy: chloroplasts are not inherited, their maintenance requires specialized host physiology, and must ultimately be replaced through renewed algal feeding.

The rarity of such mixed strategies reflects more than a lack of evolutionary opportunity. As Vermeij argues, apparently “forbidden” phenotypes often arise from functional incompatibilities and trade-offs imposed by the limited time-energy budget of organisms [42, 43]. Combining trophic modes requires the construction, regulation, and maintenance of two distinct metabolic systems, while the benefits of each may be realized under different ecological conditions [44]. Photosynthetic symbioses impose additional difficulties: the host must expose sufficient tissue to light, regulate symbiont abundance and division, induce the transfer of photosynthate, and balance competition for limiting nutrients. Moreover, photosynthates generally provide an incomplete diet, so conventional feeding is rarely abandoned [45]. In multicellular hosts, the restriction of symbionts or captured chloroplasts to particular cell types also limits their integration and transmission. Mixed trophic strategies are therefore most likely to persist when the complementary benefit is large and predictable enough to offset these coordination costs. Their rarity illustrates how energetic, developmental, and ecological constraints can leave seemingly viable regions of phenotypic space largely unoccupied.

The contrast between plants and animals is fundamentally an evolutionary problem. The relevant question is not whether plants lack cognition, but under what ecological conditions selection favors costly investments in sensing, information processing, and locomotion rather than acquisition of sessile resources based on (local) growth and physiological plasticity. This question is directly related to the long tradition of *foraging theory*, which asks how organisms allocate time and energy to locate, select, and exploit resources in heterogeneous environments [46– 51]. In animals, foraging links movement and sensory information to encounter rates and energetic returns, and has been proposed as an important selective context for the evolution of goal-directed cognition [52]. However, foraging does not necessarily require locomotion of the whole organism. Plant ecologists have long used the same concept to describe the plastic deployment of roots, shoots, and ramets toward favorable resource patches: plants explore heterogeneous environments primarily by differential growth and modular construction rather than by displacement of the individual [30, 32, 53, 54]. Resource competition theory further predicts that no single strategy can maximize efficiency on multiple limiting resources simultaneously [43, 55]. Animals and plants can therefore be viewed as solving a common resource-acquisition problem through contrasting spatial strategies—mobile search supported by sensing and information processing versus growth-mediated exploration and distributed uptake. This trade-off makes the concept of *evolutionary branching* particularly relevant. In Adaptive Dynamics, ecological feedbacks can generate disruptive selection that splits an initially monomorphic population into divergent coexisting strategies [56–62]. Evolutionary branching thus provides a natural mechanism by which continuous phenotypic variation can give rise to sharply separated ecological strategies, with intermediate forms selected against. The structure of the resources can play a central role in this process, as illustrated by resource-driven branching in other major transitions of biological organization [63]. In the present context, active sensing and locomotion should be favored when valuable resources are spatially and temporally uncertain and must be actively located, whereas their energetic costs become unnecessary when energy is predictably available and can be exploited without displacement.

Here, we formalize this trade-off using Adaptive Dynamics [56, 58, 64, 65] and spatially explicit individual-based models [66–68] in environments containing both a stable energy source (light) and patchily distributed, fluctuating resources. Starting from intermediate generalists able to exploit both resource classes, evolution repeatedly produces two divergent lineages corresponding to contrasting foraging modes: sessile, light-dependent agents that progressively abandon costly sensing and locomotion, and mobile agents that forgo light harvesting while investing in sensing and active search. Plant-like and animal-like strategies, therefore, emerge rather than being imposed by the model. The resulting scarcity of intermediate phenotypes in the linear- and convex-return cases suggests that the apparent gap between these modes of life can arise from disruptive selection acting on the energetic and ecological costs of alternative ways of finding and acquiring resources; concave returns can instead favor intermediate strategies (SI IV). In this view, the plant-animal contrast does not reflect a distinction between organisms that forage and organisms that do not, but between fundamentally different solutions to the foraging problem: one based primarily on distributed growth and persistent resource capture, the other on locomotion, sensing, and active search. Evolutionary branching can drive populations away from intermediate compromises and toward these contrasting, mutually reinforcing forms of organization.

## II. METHODS

Our main framework is an evolutionary dynamical system in which a set of agents {*A*_*k*_} exists, perishes, and evolves on a two-dimensional lattice Ω (see Figure 1). This lattice is defined as an *N*_*x*_ × *N*_*y*_ array of sites (or patches) that can contain both agents and resources.

**FIG. 1.**
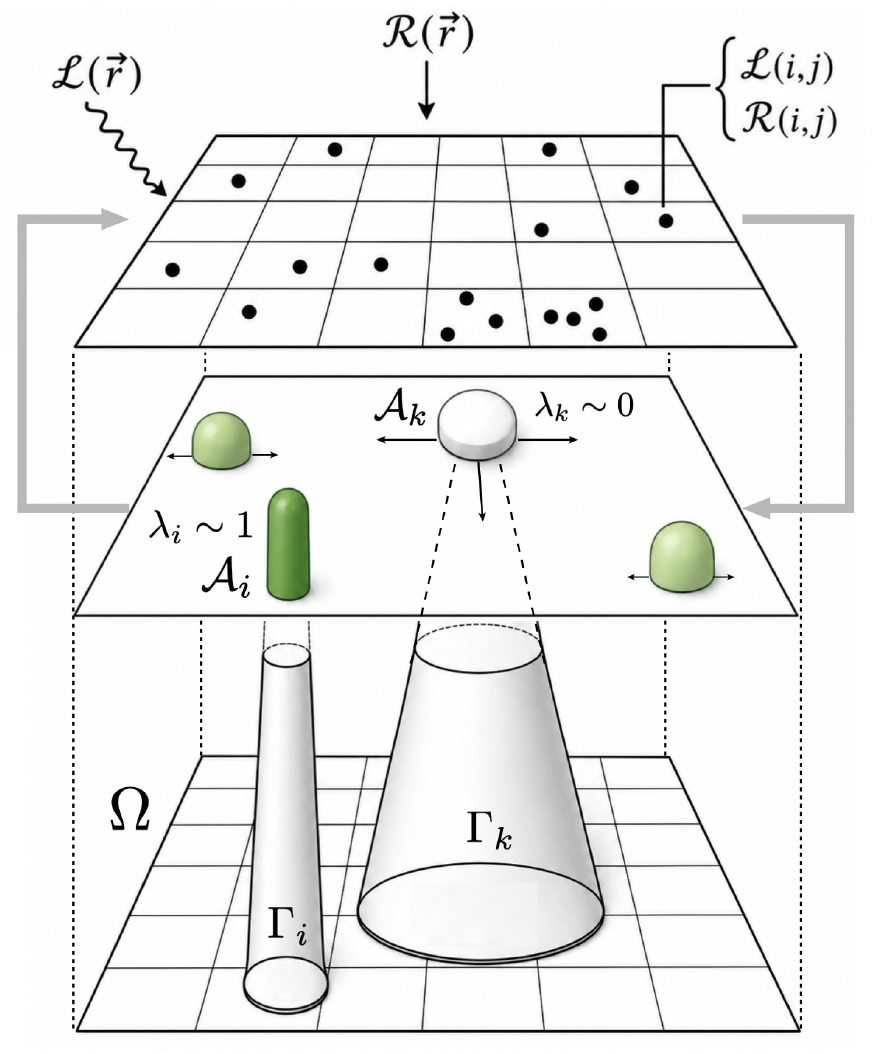
Three-layer representation of the agent-based model for the evolution of animal-like and plant-like strategies. The upper layer represents the spatial domain Ω, where light, 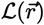, and stochastic, discrete resources, 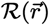 act as inputs. These fields define local resource availability at each lattice site, ℒ (i, j) and ℛ (i, j). The middle layer contains agents *A*_*k*_ located on the same domain Ω. Agents differ in the resources they exploit and in their degree of mobility, ranging from freely moving, animal-like agents shown in white to sessile, plant-like agents shown in green. The lower layer depicts the corresponding perception and movement domains available to each agent. These domains, denoted Γ_*k*_, define the spatial range over which agents can detect, access, or move toward resources, thereby linking energetic strategies with movement constraints. A parameter 0 ≤ λ ≤ 1 associated with each agent (and evolving through the simulation) weights the relative use of light (λ ∼ 1) or material resources (λ ∼ 0).

Specifically,

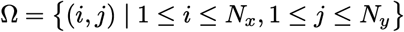

with periodic boundary conditions.

### A. Resources

Each patch (*i, j*) contains two distinct resources: a non-depletable resource ℒ(*i, j*) and a depletable resource ℛ(*i, j, t*). The non-depletable resource ℒ (*i, j*) models light irradiation, represented as a fixed, spatially dependent quantity across the domain Ω. We examine two spatial distributions for ℒ (*i, j*): a homogeneous configuration, where ℒ (*i, j*) = ℒ is constant across the entire domain, and a non-homogeneous configuration, where irradiation decreases linearly along one spatial dimension according to the gradient

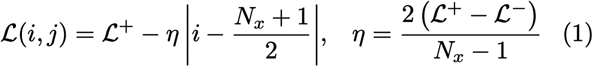

where (ℒ^+^, ℒ^−^) are the maximum and minimum irradiation levels, and *η* is the gradient slope.

The discrete and depletable resource ℛ(*i, j, t*) signifies the number of discrete particles at the site (*i, j*) at time *t*. Its dynamics are controlled by local, spatially independent, and memoryless stochastic processes for degradation and replenishment. The degradation function

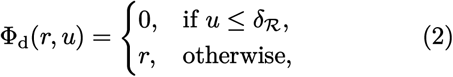

captures the decay of the resource, effectively reducing the resource *r* to zero with probability *δ*_R_, in a density-independent manner.

The replenishment function Φ_*r*_ introduces new particles independently of the current resource level. The number of incoming particles at each site follows a geometric distribution with mean *α*. This is generated from a uniform variate *u*_r_ via the inverse transform method,

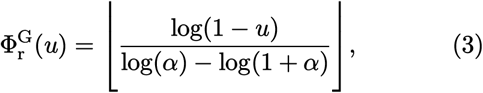

which satisfies, for all *k* = 0, 1, 2…,

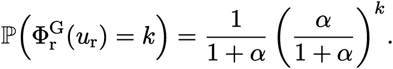

This form is used to generate uncertainty in the environment.

The combined update from time *t* to *t* + 1 is given by the composite function

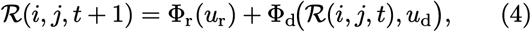

where *u*_d_, *u*_r_ ∼ *U*(0, 1) are independent random numbers drawn for each site and time step. In addition, ℛ can be depleted by agents, which will be elaborated in due course.

### B. Agent Description

An agent is characterized by a tuple

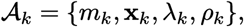

where *m*_*k*_ ≥ 0 denotes the agent’s mass, and **x**_*k*_ ∈ Ω its spatial location. The variable *λ*_*k*_ ∈ [0, 1] represents the fraction of the agent’s normalized energy budget allocated to exploiting the non-depletable resource (light); consequently, 1 − *λ*_*k*_ is the fraction directed toward harvesting the depletable resource ℛ. The agent’s sensing capability for **ℛ** is governed by *ρ*_*k*_ ∈ {0, 1, …, *ρ*_max_}, which defines an effective search radius *r*_*k*_, defined only for *ρ*_*k*_ *>* 0, *r*_*k*_ := *ρ*_*k*_ −1. If *ρ*_*k*_ = 0, the agent is completely oblivious to its environment.

Let Γ_*k*_ denote the set of lattice sites within this neighborhood

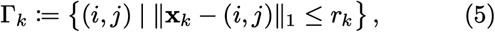

where ∥ · ∥_1_ is the Manhattan distance. The corresponding set of observable resource patches is

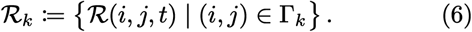

The agent assesses and can reach all patches in Γ_*k*_, but can only forage from the patch it currently occupies.

### C. Agent Evolution Rules

At a time step *t*, all agents are asynchronously updated, in a randomly selected order, each step following this outline; the complete pseudocode is given in SI 1 and 2. For an agent *A*_*k*_, the step is as follows. With probability *δ*_*A*_, the agent randomly dies. If the agent survives, then resource gathering and potential movement follow.

For agents with *ρ*_*k*_ *>* 1, acquisition of the depletable resource ℛ drives movement; an agent with *ρ*_*k*_ = 1 perceives only its own site and therefore remains stationary.

The agent evaluates the legal candidate set

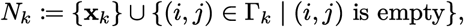

which includes the current site and excludes occupied destinations, computing the net foraging gain *F*_*k*_(*i, j*) for each candidate:

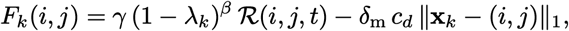

where *γ* denotes the resource-to-mass conversion parameter, *c*_*d*_ *δ*_m_ is the cost of moving one lattice unit, and *β >* 0 is the returns exponent acting on the energy-allocation budget. The agent chooses a maximizer,

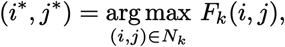

and moves to this location only if the gain is strictly positive; if several sites tie for the maximum, one is selected uniformly at random. The movement distance *d*_*k*_ is thus

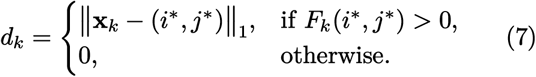

After any movement, the agent’s position is updated to **x**_*k*_ = (*i*^*^, *j*^*^), or remains unchanged if *d*_*k*_ = 0.

The agent updates its mass based on resource intake and metabolic losses. At its updated location (*i, j*), the agent gains mass by harvesting the non-depletable light resource ℒ(*i, j*) and, when capable of foraging, by consuming the depletable resource ℛ(*i, j, t*). These contributions are combined into a gross mass-gain function,

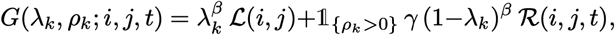

where *γ* is the efficiency with which acquired resources are converted into biomass and *β >* 0 is the returns exponent on the allocation budget. By default, constant marginal returns *β* = 1 is assumed, for which the gross returns are linear in the allocation.

The agent also loses mass due to phenotype-dependent maintenance. The normalized maintenance cost is defined as a linear combination of costs,

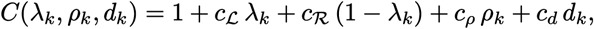

where the coefficients *c*_ℒ_, *c*_ℛ_, *c*_*ρ*_, and *c*_*d*_ represent additional maintenance losses associated with light harvesting, substrate processing, foraging, and movement, respectively, all expressed relative to a unitary, phenotype-independent baseline loss. The net mass change over the time step is the difference

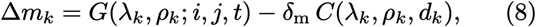

where *δ*_m_ is a scaling parameter that sets the magnitude of mass loss. If, after the update, the agent’s mass becomes non-positive—that is, if *m*_*k*_ + Δ*m*_*k*_ ≤ 0—the agent perishes due to starvation. Otherwise, its mass is updated and clamped to the plausible range *m*_*k*_ ∈ [0, 1]:

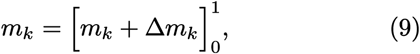

where 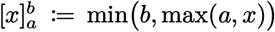. Reproduction is possible only if the agent’s mass before clamping meets or exceeds the unit threshold,

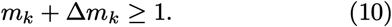

In this case, a neighboring lattice site is selected uniformly at random. If the selected site is already occupied, reproduction is aborted, and the update for this agent then concludes. If that site is unoccupied, the agent reproduces by producing an offspring *A*_*l*_ at the chosen location. Upon reproduction, the parent’s mass is halved, and the offspring is initialized with the same reduced mass.

The offspring inherits the parent’s traits *λ*_*k*_ and *ρ*_*k*_, which may then undergo mutation. With probability *µ*, the energy allocation parameter is perturbed by Gaussian noise:

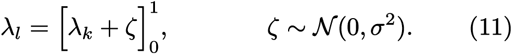

Independently, with probability *µ*, the sensing capability is mutated by a unit step in either direction with equal likelihood. Let *ε* be a Rademacher random variable satisfying ℙ(*ε* = 1) = ℙ(*ε* = −1) = 1*/*2. The mutated value is obtained by adding *ε* to the parent’s sensing capability and clamping the result to the admissible range

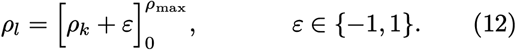

## III. RESULTS

### A. Branching on a Slope

Introducing a spatial light gradient (Eq. (1)) transforms environmental heterogeneity into a direct driver of phenotypic diversification [69]; the precise periodic implementation and its interpretation are given in SI I.A. High-irradiation regions select for autotrophic investment (*λ* → 1), while low-irradiation peripheries favor heterotrophic foraging (*ρ >* 0).

Figure 2 presents three representative realizations across distinct gradient slopes, illustrating the emergent spatial organization and phenotypic composition of the population. In the case of a steep gradient, the domain partitions into three distinct ecological zones. A high-irradiation region supports an almost exclusive population of plant-like organisms, which allocate their budget to light harvesting while incurring negligible maintenance costs for sensing and movement. Conversely, the low-irradiation periphery becomes dominated by animal-like foragers exploiting available resources in areas that do not support plant growth. Between these extremes lies a transitional zone of intermediate light intensity, where neither strategy achieves complete dominance, resulting in a polymorphic coexistence maintained by frequency-dependent selection and movement-mediated spatial redistribution.

**FIG. 2.**
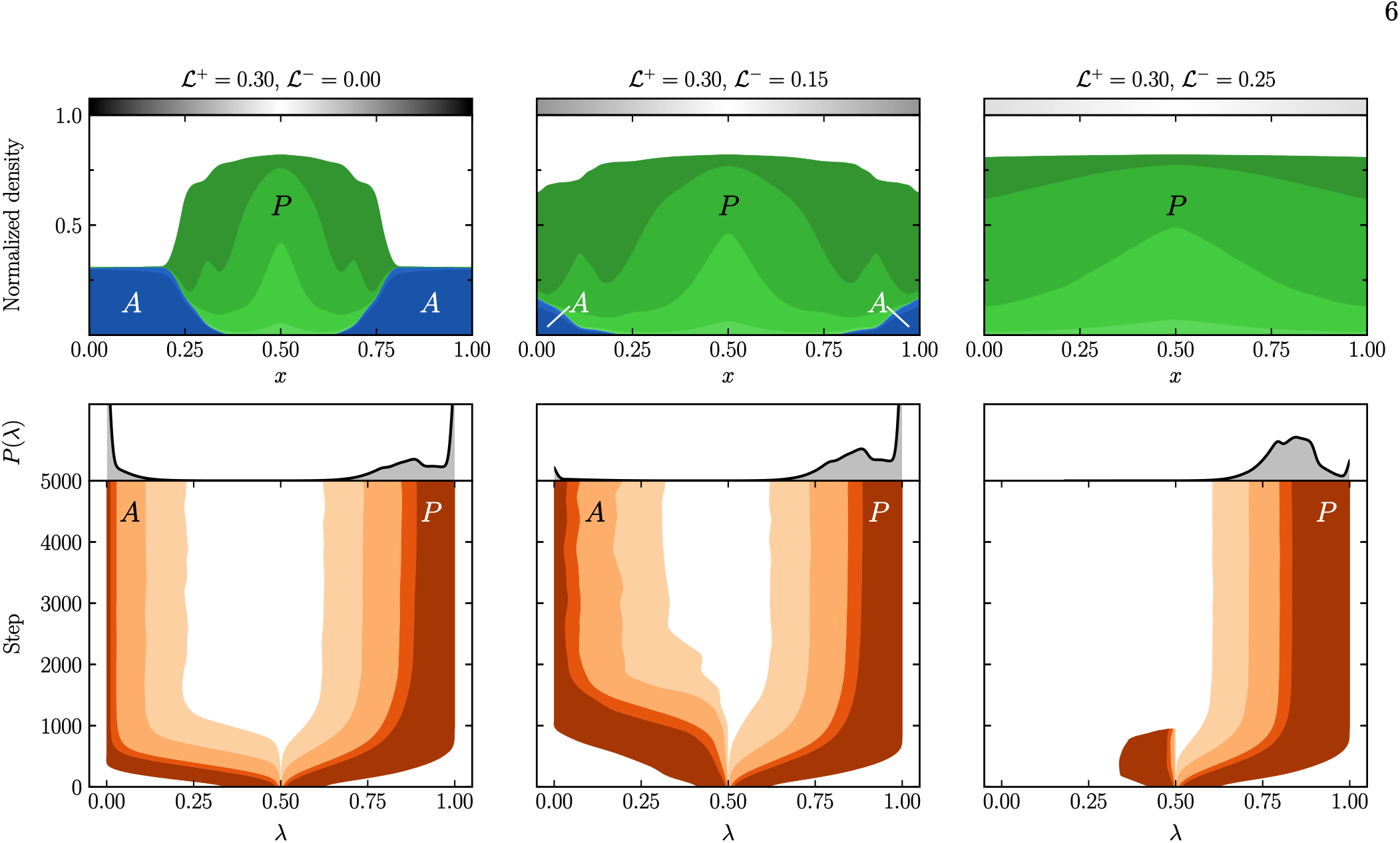
Evolutionary differentiation of plant-like and animal-like strategies along spatial light gradients. Columns correspond to three light profiles ℒ (x) (grayscale bars, top), ranging from steep (left) to shallow (right), with the rightmost approaching an almost uniform, high-illumination regime. Upper panels show final normalized spatial densities, colored by resource-allocation trait λ (animal-like foragers, λ < 0.5, blue; plant-like autotrophs, λ > 0.5, green). Lower panels show the evolutionary distribution of λ over time; shaded orange regions indicate one-sided cumulative quantile bands (50–68–95–99.7% of trait mass) integrated from each phenotypic extreme toward the mixed center, with the median trajectories corresponding to the inner boundary of the darkest band. Long-term marginal distributions P(λ) are shown above each trajectory. Under a steep spatial gradient, plants dominate the well-illuminated zone, whereas animals occupy the dark periphery, with a transitional coexistence zone between them. As the gradient becomes shallower, the spatial niche available to foragers contracts, such that plant-like agents colonize the entire domain and restrict animals to the dimmest margins. Under the almost uniform, high-light condition, the forager branch initially emerges but is subsequently outcompeted by lineages evolving toward higher λ; nevertheless, the global surplus of light and material resources sustains intermediate mixotrophic strategies—agents that primarily harvest light yet retain sufficient foraging capacity to opportunistically exploit rare, high-yield patches—thereby preventing full convergence to the pure plant phenotype. Parameters: α = 0.5, β = 1, γ = 0.1, δ_*A*_ = 0.1, δ_ℛ_ = 0.1, δ_m_ = 10^−2^, µ = 0.1, ρ_max_ = 5, σ_*λ*_ = 0.05, c_*d*_ = 1, c_ℒ_ = 1, c_ℛ_ = 1, c_*ρ*_ = 1. Light levels ℒ^+^ and ℒ^−^ vary by column as indicated by the grayscale bars.

A shallower gradient contracts the spatial niche available to foragers. If light remains sufficient for autotrophic strategies across most of the domain, plant-like agents colonize the entire domain. However, foragers may persist at the dimly lit periphery, where reduced light lowers plant density and leaves resources underexploited. This creates a narrower but stable niche where animals coexist with the surrounding plant population, sustained by their superior ability to locate and exploit residual unoccupied resource patches.

Under conditions of uniformly high illumination, the forager branch initially emerges but is subsequently out-competed by lineages branching toward higher *λ*. Nevertheless, the global surplus of light and abundance of resources allow for the persistence of intermediate, mixotrophic strategies. These agents primarily harvest light but retain sufficient sensing and foraging capabilities to opportunistically exploit rare, high-yield resource patches. This allows them to subsist on reliable harvesting returns from light while also reproducing much faster than the pure plant phenotype when resourcedense patches appear within their sensing range.

Spatial variation in selection, combined with mobility, thus translates environmental heterogeneity into phenotypic differentiation: local resource regimes select for autotrophs or heterotrophs, while the steepness of the gradient determines the width of the transitional coexistence zone.

#### Nonlinear returns

The persistence of intermediate strategies observed under uniform high illumination (Figure 2) depends on the assumption of linear returns on investment (*β* = 1); the corresponding concave- and convex-return analysis is given in SI IV. When allocation traits map linearly to energetic intake, partial investment in both light harvesting and material foraging yields proportional benefits, permitting mixotrophs to exploit dual niches without severe efficiency penalties. However, in natural systems, resource acquisition and processing pathways typically exhibit nonlinear scaling across ecologically relevant investment ranges.

Biological mechanisms underlying resource capture can produce convex gain functions within operational regimes. Photosynthetic apparatuses require substantial fixed investments in pigment-protein complexes, electron transport chains, and carbon-fixation enzymes before achieving functional throughput. Similarly, heterotrophic foraging involves fixed cognitive and locomotor overheads: sensory organs must reach minimum complexity to resolve environmental cues, and motor systems require baseline sophistication. Partial investment in either system thus incurs disproportionate costs relative to returns, rendering generalist strategies energetically suboptimal compared to specialists that concentrate resources above minimally functional thresholds.

Figure 3 shows how superlinear returns reshape trait distributions across gradient steepnesses. Under linear returns (*β* = 1), trait distributions retain significant mass in intermediate trait allocation. As *β* increases above unity, probability density shifts toward the phenotypic boundaries (*λ* → 0 and *λ* → 1), with corresponding depletion in the central mixotrophic zone (Figure 3 b, e). Median trajectories (Figure 3 c, f) confirm that populations converge to more specialized strategies under increasing returns.

**FIG. 3.**
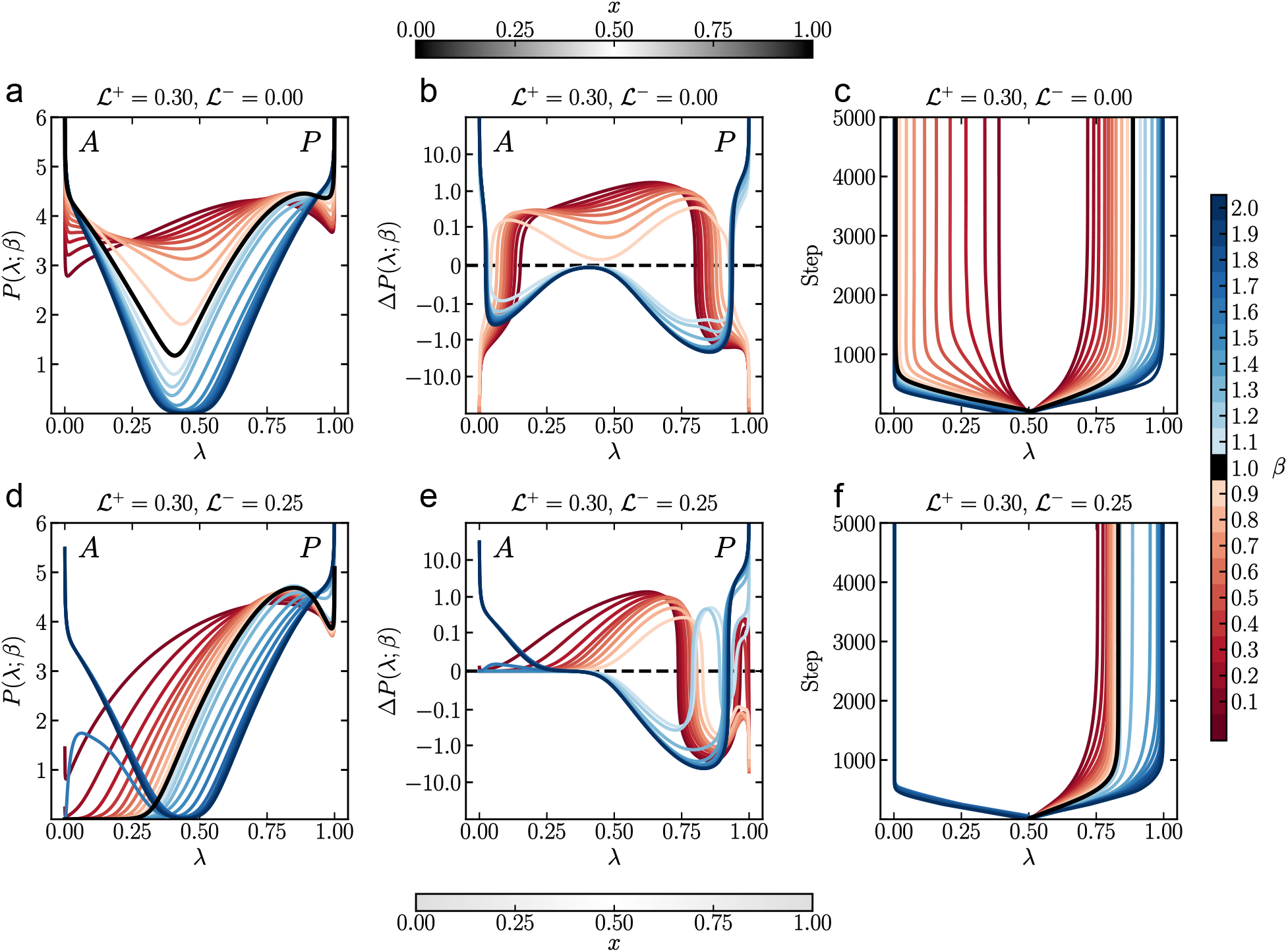
Phenotypic distributions under varying return-on-investment regimes. The top and bottom rows depict steep (ℒ^−^ = 0.0) and shallow (ℒ^−^ = 0.25) light gradients, respectively, with ℒ^+^ = 0.30 held constant in both cases. Panels (a) and (d) illustrate the pseudo-logarithmically transformed long-term marginal probability, P(λ; β), of the energy allocation trait λ. Curves are color-coded according to the returns exponent β using a diverging colormap. The linear reference case (β = 1) is depicted in black; red denotes diminishing returns β ∈ (0, 1), which favor mixotrophy, whereas blue indicates superlinear returns β ∈ (1, 2], which favor specialization. Panels (b) and (e) display the pseudo-logarithmically transformed differences in marginal probabilities relative to the linear reference, defined as ΔP (λ) = P(λ; β) − P(λ; β = 1). Positive values indicate phenotypic conditions favored by selection, while negative values denote phenotypes selected against, relative to the linear baseline. Panels (c) and (f) illustrate the evolutionary trajectory of the median λ for the animal-like (λ ≤ 0.5) and plant-like (λ ≥ 0.5) subpopulations. In all cases, sublinear returns (β < 1) select mixotrophic strategies and shift the medians toward the center, whereas superlinear returns (β > 1) select for specialization and shift the medians toward the extremes. Parameters: α = 0.5, γ = 0.1, δ_*A*_ = 0.1, δ_ℛ_ = 0.1, δ_m_ = 10^−2^, µ = 0.1, ρ_max_ = 5, σ_*λ*_ = 0.05, c_*d*_ = 1, c_ℒ_ = 1, c_ℛ_ = 1, c_*ρ*_ = 1.

Convex returns (*β >* 1) can also drive a fundamental restructuring of community composition. This is illustrated in the shallow gradient scenario, where intensified selection for extreme traits drives mixotrophs to extinction. Because mixotrophs—unlike plants—compete with foragers for shared resources, their disappearance alters local resource dynamics. This release from competitive pressure allows resources to accumulate in formerly mixotroph-dominated patches, opening a niche for foragers and enabling the evolution of the animal branch.

### B. Branching in Homogeneous Environments

In contrast to the spatially heterogeneous scenarios examined previously, we now consider the case of uniform light distribution across the domain, where ℒ(*i, j*) = ℒ for all (*i, j*) ∈ Ω. This homogeneous configuration eliminates spatial variation, thereby removing the geographic niche partitioning mechanism that primarily drove diversification in the gradient case. Nevertheless, phenotypic diversification persists through frequency-dependent selection mediated by competition for finite space and the stochastic, depletable resource ℛ.

Figure 4 presents a comprehensive phase diagram characterizing the long-term evolutionary outcomes as a function of ambient light intensity ℒ. Panel (a) displays the joint probability density *P* (*λ*, ℒ), revealing how the distribution of the energy-allocation trait *λ* shifts across environmental conditions. Panels (b)–(e) provide columnar cross-sections of the joint trait distribution *P* (*λ, ρ*) at specific light levels, illustrating the trade-off between light harvesting (*λ*) and spatial sensing/foraging capability (*ρ*).

**FIG. 4.**
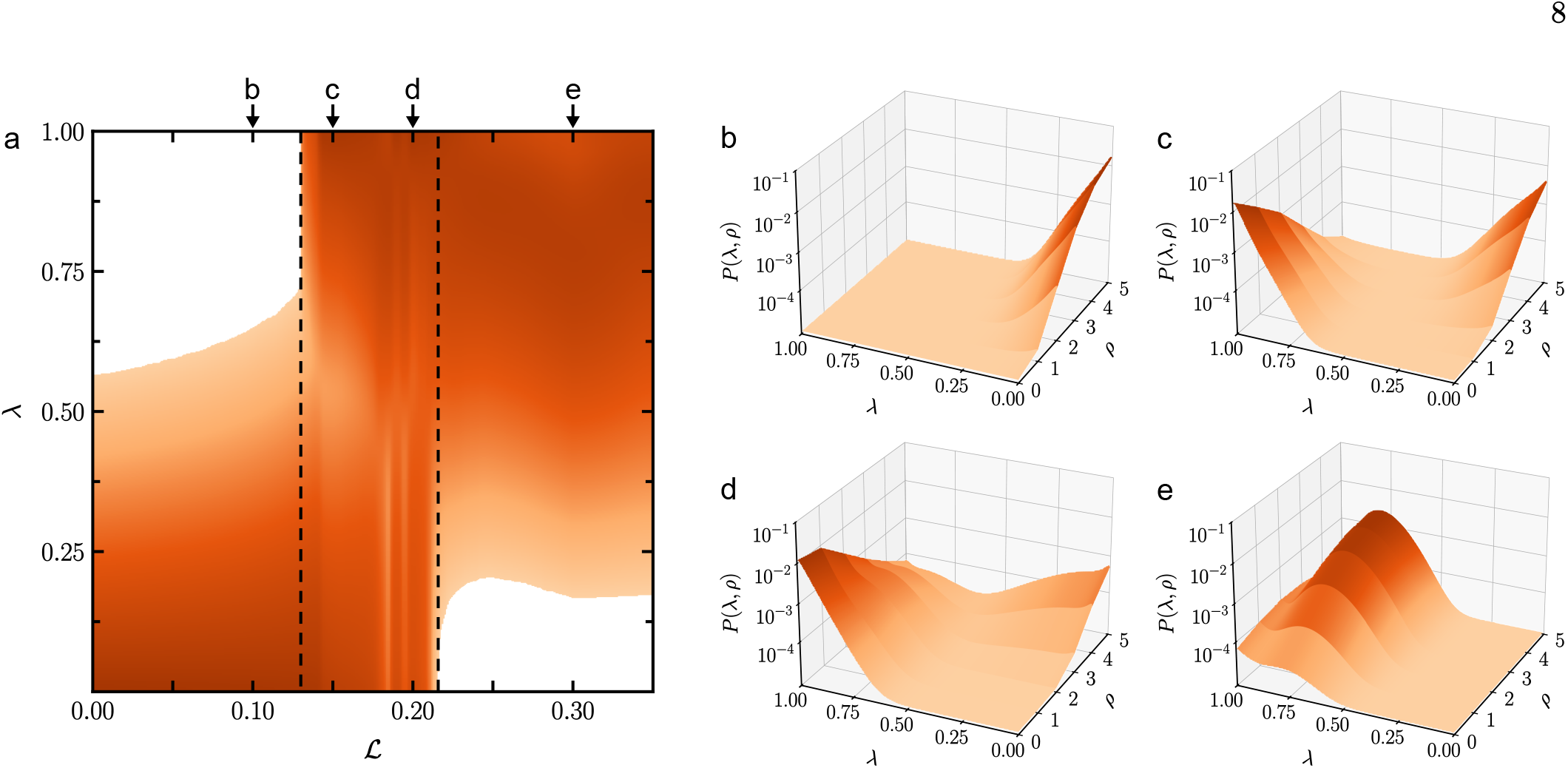
Evolution of phenotypic distributions across varying light availability in homogeneous environments. Panel (a) displays the long-term normalized density distribution of the light-harvesting trait λ as a function of uniform environmental light intensity ℒ. The color mapping indicates the marginal probability *P*(λ) on a logarithmic scale, highlighting regions of high phenotypic abundance. Dashed lines delineate the coexistence regime where animal-like and plant-like phenotypes co-occur. Panels (b–e) present cross-sections of the joint probability density *P*(λ, ρ) at selected light levels. In panel (b), at low light ℒ = 0.1, the population is dominated by animals (low λ, high ρ), as scarce light energy cannot support mostly phototrophic strategies or the intermediate allocations leading to plants. In panel (c), as light rises to ℒ = 0.15, a bimodal distribution emerges, with stable coexistence of animals (low λ, high ρ) and plants (high λ, low ρ). In panel (d), at increased light ℒ = 0.2, the plant peak strengthens leaving much less available space for animals even if resources remain abundant. In panel (e), under light-saturated conditions ℒ = 0.3, the fitness landscape flattens, weakening selective pressures for strict specialization. The dominant phenotype shifts toward organisms that rely mainly on light harvesting but retain some foraging ability (ρ > 0) to maximize division rates by opportunistically feeding on nearby resource-rich patches. Parameters: α = 0.5, β = 1, γ = 0.1, δ_*A*_ = 0.1, δ_ℛ_ = 0.1, δ_m_ = 10^−2^, µ = 0.1, ρ_max_ = 5, σ_*λ*_ = 0.05, c_*d*_ = 1, c_ℒ_ = 1, c_ℛ_ = 1, c_*ρ*_ = 1.

When light intensity is low, autotrophic investment yields insufficient energy to sustain population growth. In this regime, selection strongly favors agents with low *λ* values and non-zero sensing radii (Figure 4b). These “animal-like” heterotrophic specialists exploit spatial and temporal fluctuations in ℛ through active foraging, a strategy that remains viable independently of light levels.

As light availability increases, the system enters a coexistence regime delimited by dashed lines in Figure 4a. Here, frequency-dependent selection produces a bimodal distribution of phenotypes. For the linear-return model, the SI analysis identifies this as saddle-driven two-optima divergence rather than classical convergence-stable branching (SI IV.D). As illustrated in Figures 4c and d, two distinct peaks emerge in the *λ*-*ρ* landscape: a cluster at high *λ* and *ρ* = 0 and one cluster at low *λ* and high *ρ*, corresponding to plant-like and animal-like strategies, respectively. This polymorphism is stable because in this parameter regime, each specialist out-performs generalists in its respective niche; intermediate “mixotrophic” strategies pay the maintenance costs for both systems without achieving competitive efficiency in either, leading to their exclusion via the “curse of mixed strategies.”

At sufficiently high light intensities, the selective pressure favoring strict specialization weakens. As shown in Figure 4e, the dominant phenotype shifts toward high *λ* but retains non-zero *ρ*. In this light-saturated regime, the marginal benefit of additional light harvesting diminishes, while the ability to sense and move toward resource-rich patches provides a supplementary advantage for maximizing division rates. Consequently, the population converges on a single peak characterized by high autotrophic investment coupled with moderate foraging capability, reflecting a relaxation of the stricter trade-off observed at lower light levels.

Thus, such phenotypic differentiation does not require spatial heterogeneity. In a homogeneous environment, the intrinsic trade-offs between the two resource-acquisition strategies alone generate frequency-dependent selection. The mechanism transitions from direct competition-mediated coexistence at intermediate light levels to mutation-driven exploration of trait space under light-saturated conditions.

### C. Adaptive Dynamics

To complement the agent-based model, we formulate a trait-based Adaptive Dynamics (AD) model of the population’s long-term evolution in a continuous phenotypic space [57, 69–72]. This is a separate, simplified well-mixed model that asks whether analogous diversification can arise from physiological trade-offs alone. The AD model replaces local stochastic feedbacks and explicit movement costs with deterministic global resource regulation and a saturating uptake response; its detailed analysis is given in SI IV. Within the AD framework, a large, well-mixed population is assumed, and spatial heterogeneity and demographic stochasticity are neglected [71]. Its parameters are model-specific and need not match those of the ABM, and its results do not constitute a derivation of the ABM dynamics.

The phenotypic composition is represented by a continuous density function *n*(*λ, ρ, t*), where *λ* ∈ [0, 1] denotes the proportion of resources allocated to light harvesting (autotrophy) and *ρ* ≥ 0 represents the foraging radius, treated as a continuous nonnegative trait. Mutations are assumed to be rare and to induce only small phenotypic deviations; consequently, the population remains effectively monomorphic—characterized by a single resident phenotype (*λ, ρ*)—except during transient invasion events triggered by rare mutants. The ecological feedback between individuals and their environment is captured through a system of two coupled ordinary differential equations governing the dynamics of the resource concentration ℛ(*t*) and the total population density *N* (*t*), the integral of the population density *n*(*λ, ρ, t*) over all admissible trait combinations:

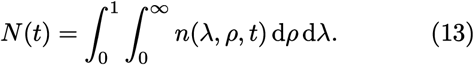

Following the agent-based mass update above, we collect all maintenance-related losses into a single normalized cost function,

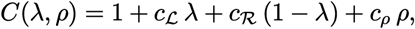

where the coefficients *c*_ℒ_, *c*_ℛ_, and *c*_*ρ*_ are the additional maintenance losses associated with light harvesting, substrate processing, and foraging, respectively, expressed relative to a unitary baseline loss. Individual fitness is defined by the per-capita growth rate function *f* (*λ, ρ*; ℛ), which depends on the physiological traits (*λ, ρ*), the environmental resource concentration ℛ, and constant ambient light ℒ. It can be written as

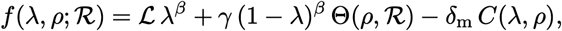

where *γ* denotes the efficiency with which acquired resources are converted into biomass, *δ*_m_ scales the cost function, and *β* controls how much investment affects returns. Most of the presented results are derived from the linear returns case, *β* = 1, unless explicitly stated. A more in-depth exploration of the concave and convex returns regimes is presented in SI IV.

Resource uptake follows a saturating Monod (Holling type II) functional response [73, 74],

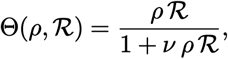

where the handling time *v* imposes diminishing returns on foraging efficiency, preventing unbounded gains and thereby ensuring well-behaved evolutionary dynamics.

The dynamics of forageable resources are modeled according to chemostat-like principles [75],

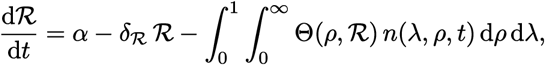

with *α* representing the constant external input rate of the resource and *δ*_ℛ_ the combined rate of abiotic decay and dilution. The integral term captures the total consumption by the population across all trait values. Population dynamics incorporate density-dependent regulation through a logistic-type term with an explicit carrying capacity *K* in a homogeneous environment,

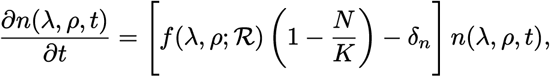

where *K* is the carrying capacity of the homogeneous AD model, and *δ*_*n*_ is its background mortality rate. These parameters are model-specific and need not equal the corresponding quantities in the ABM.

Within the Adaptive Dynamics framework, evolutionary change is governed by the sign of the invasion fitness *σ*(*λ*_m_, *ρ*_m_; *λ, ρ*), defined as the initial per-capita growth rate of a rare mutant phenotype (*λ*_m_, *ρ*_m_) introduced into a resident population at its ecological equilibrium,

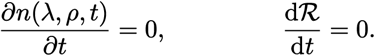

At this equilibrium, the invasion fitness simplifies to

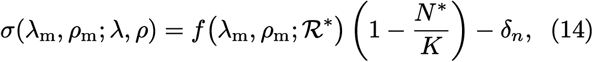

where ℛ^*^ and *N* ^*^ are the equilibrium resource level and resident population density, respectively, determined by the resident phenotype (*λ, ρ*).

Evolution proceeds through incremental mutational steps, with the direction and rate of phenotypic change dictated by the selection gradient—the partial derivatives of invasion fitness with respect to mutant traits— evaluated at the resident phenotype [71],

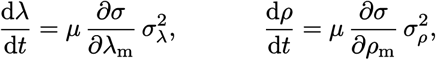

where *µ* is the mutation rate and 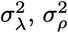 denote the mutational variances of the two traits (the corresponding standard deviations *σ*_*λ*_, *σ*_*ρ*_ are reported in Figure 5). Evolutionary singular strategies (*λ*^*^, *ρ*^*^) are defined by the vanishing of the

**FIG. 5.**
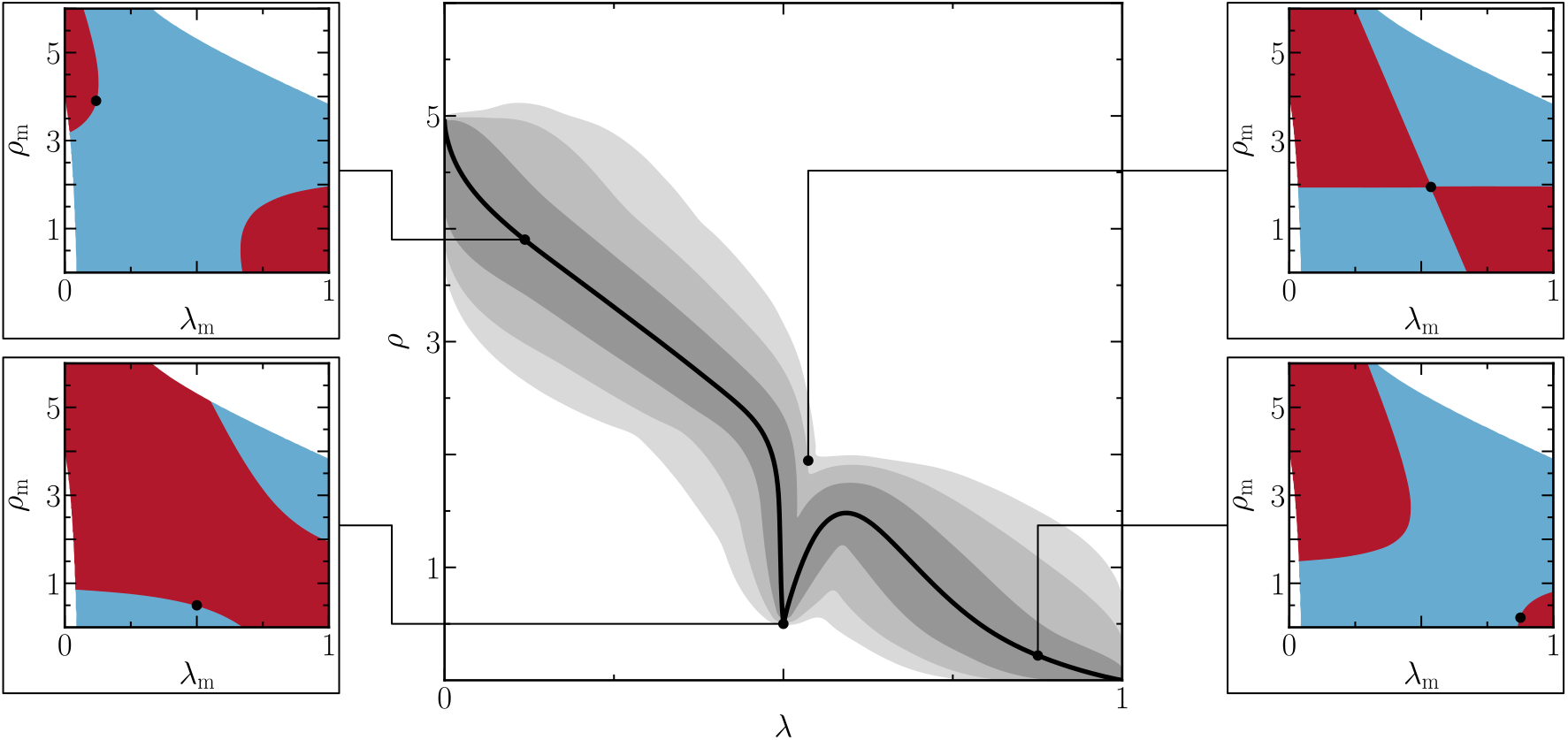
Evolutionary trajectories of the traits λ and ρ under Adaptive Dynamics. Solid lines denote the ensemble expectation, with surrounding shaded regions representing the central statistical spreads corresponding to 1-3 σ (encompassing approximately 68-95-99.7% of realizations). The middle plot captures a bifurcation event in which stochastic evolutionary dynamics split into two distinct lineages. All trajectories start from an initial resident population with traits (λ, ρ) = (0.5, 0.5), and trait evolution is driven by successful invasions stemming from normally distributed mutations characterized by standard deviations σ_*λ*_ = 10^−2^ and σ_*ρ*_ = 10^−1^. Insets display the invasibility plot for resident populations in the expected evolutionary trajectory and the interior singular strategy: dark red regions indicate mutant traits capable of invading the resident population; light blue regions correspond to non-invasible mutants; blank areas denote trait combinations that cannot sustain a viable population. Parameters: ℒ = 1, c_ℒ_ = 0.5, γ = 5, *v* = 1, c_ℛ_ = 1, c_*ρ*_ = 0.25, δ_m_ = 10^−1^, α = 10^−1^, δ_ℛ_ = 10^−1^, δ_*n*_ = 10^−2^, µ = 10^−2^, β = 1.

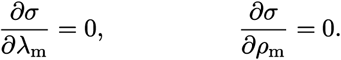

Higher-order derivatives of the invasion fitness determine the evolutionary fate of such singular points; the full Hessian, convergence-stability, and boundary-equilibrium analysis is provided in SI IV and IV.D. Let *H* denote the Hessian matrix of second partial derivatives of *σ* with respect to (*λ, ρ*) evaluated at the singularity,

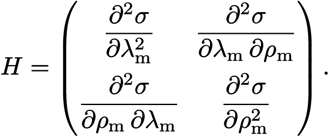

When *H* is negative definite, the singularity constitutes an evolutionarily stable strategy (ESS), meaning nearby mutant phenotypes are unable to invade it [76, 77]. If, in addition, the selection gradient directs trait trajectories toward the singularity from neighboring regions of trait space, the strategy is convergence stable [78]. A singular strategy that is convergence stable but not evolutionarily stable is a branching point: disruptive selection drives the population toward phenotypic divergence and the emergence of distinct types [78]. At the linear-return exponent *β* = 1 adopted here, however, this classical scenario is not realized: the divergence between sessile and mobile strategies is instead the two-optima mechanism described below.

At the linear-return exponent *β* = 1 adopted in the simulations, the second derivative with respect to *λ*_m_ vanishes identically because the fitness function is linear in the allocation trait,

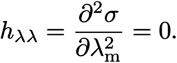

The cross-derivative is obtained by differentiating the marginal foraging benefit with respect to *λ*_m_,

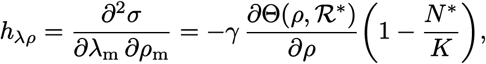

which is negative because the conversion efficiency *γ*, the marginal resource uptake *∂*_*ρ*_Θ, and the crowding factor are all positive; this encodes the physiological trade-off whereby increased autotrophic allocation diminishes the marginal fitness returns of foraging. The remaining entry is negative as well,

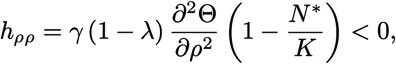

since the Monod response is concave in *ρ*. Consequently, the Hessian determinant simplifies to

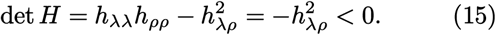

This negative determinant confirms that the Hessian is indefinite and that the singular strategy is a saddle point rather than an ESS.

Different phenomena are commonly called branching. Classical (Geritz) branching [78] needs an interior singular point that is convergence stable but not evolutionarily stable: the population converges to a generalist and is then split. In the linear-return case used here (*β* = 1), no such point exists: the interior is a repelling saddle of the selection gradient (the Jacobian *J* of the resident selection gradient has det *J <* 0), so a lineage never converges to the generalist. The split comes instead from frequency-dependent selection between the two boundary optima, the plant (*λ* → 1, *ρ* → 0) and the animal (*λ* → 0, *ρ >* 0). Each specialist creates the environment in which the other is favored: plants leave the particulate resource unexploited, where foraging pays; foragers deplete it and compete with each other, leaving space empty for autotrophs to thrive. Thus, each specialist can invade the environment created by the other, yielding a protected dimorphism. For *β >* 1 the returns are convex and the pressure against intermediates is stronger, but the mechanism driving the “branching” is the same.

Simulations of the continuous-trait AD model illustrate this process. Starting from a generalist mixotroph at ecological equilibrium, the reported stochastic realizations end in two coexisting specialist lineages, as shown in Figure 5: (i) a plant-like strategy specializing in light harvesting with minimal foraging investment, and (ii) an animal-like strategy abandoning autotrophy in favor of active resource acquisition. The invasibility plots provide a geometric illustration of the analysis. At the singular strategy, the local invasion-fitness landscape exhibits a saddle topology, with a locally disruptive direction associated with the autotrophy–foraging trade-off. Consequently, mutants that shift allocation along the autotrophy–foraging axis toward either extreme have positive invasion fitness, as they are able to exploit underutilized niches created by frequency-dependent competition.

This phenotypic divergence arises directly from the eco-evolutionary feedback embedded in our model. The fitness function provides several constraints: (i) a fixed allocation budget between autotrophic and heterotrophic functions, (ii) diminishing returns on foraging due to fixed handling time *v*, (iii) trait-specific maintenance costs (*c*_L_, *c*_R_, *c*_*ρ*_), and (iv) a uniform baseline metabolic maintenance *δ*_m_ paid by all phenotypes, which does not affect the trait trade-offs. Together, these generate negative frequency-dependent selection: as the population consumes shared particulate resources, it depletes the very substrate that heterotrophs rely on, thereby favoring individuals that shift toward autotrophy—and vice versa. At intermediate phenotypes, neither strategy is optimized, creating a fitness valley that drives disruptive selection.

In this simplified, well-mixed AD setting, the analysis shows that an autotroph–heterotroph dichotomy can arise from internal life-history trade-offs alone, without requiring predation or an environmental gradient.

## IV. DISCUSSION

The contrast between plant and animal cognition is fundamentally an evolutionary problem rather than a purely mechanistic one. Cognition, movement, and sensing are metabolically costly traits that can only be maintained if environmental conditions provide sufficient fitness returns. In spatially stable and predictable environments, selection may favor sessile, low-information strategies based on growth and physiological plasticity. In contrast, when resources are patchy and fluctuate stochastically in space, active search and sensory integration can become advantageous despite their energetic costs.

By embedding these trade-offs in a computational ecological model, we allow phenotypic strategies to emerge from resource structure and energetic constraints rather than presupposing “plant-like” or “animal-like” categories. Our results reveal two distinct routes to phenotypic differentiation between sessile autotrophs and mobile heterotrophs. Under a spatial light gradient, environmental heterogeneity creates geographic niche partitioning that directly drives phenotypic divergence: plants dominate high-irradiation zones, animals occupy the periphery, and an intermediate region supports transient coexistence through movement-mediated competition. In homogeneous environments, differentiation arises instead through frequency-dependent selection alone: competition for the shared depletable resource generates disruptive selection on the allocation trait *λ*, causing a single mixotrophic lineage to split into two specialist lineages without any spatial template. The Adaptive Dynamics analysis shows how the same physiological trade-off can generate diversification without spatial structure, while also clarifying the distinction between classical branching and saddle-driven two-optima divergence.

These findings reframe the plant-animal cognition debate as an evolutionary bifurcation problem. When a resource is reliably available, as light is for autotrophs, the selective pressure for costly information-processing machinery is weak, and sessile strategies prevail through efficient conversion of predictable energy. When resources are patchy and uncertain, the benefits of movement and sensing can offset their metabolic costs, favoring mobile, cognitive agents. The coexistence of both strategies from a single mixotrophic ancestor, when it occurs under the modeled parameters, shows that an external spatial driver is not necessary.

Several limitations should be acknowledged. Our model abstracts away many biological details, including predation, social learning, memory, and the distinction between sensing and movement costs. The lattice-based spatial structure and discrete resource particles are a simplification of real ecological landscapes. Future work could explore how additional factors—such as predator-prey dynamics, environmental periodicity, spatial memory, or explicit neural costs—affect the branching conditions and the resulting cognitive strategies. Information-theoretic characterizations of environmental uncertainty also warrant further investigation as potential predictors of evolutionary transitions.

The broader implication is that the divergence of cognitive strategies need not be explained by external drivers alone; internal metabolic trade-offs can generate diversification under a wide range of environmental conditions. This suggests that the plant-animal dichotomy in cognitive complexity may be a natural outcome of eco-evolutionary dynamics rather than a taxonomic accident. By casting the question in terms of adaptive trade-offs and evolutionary diversification, our framework turns a longstanding debate about plant cognition into a quantitatively tractable problem about resource allocation, environmental statistics, and the conditions under which natural selection favors sensing and movement.

## ACKNOWLEDGMENTS

The authors thank the Complex Systems Lab members for fruitful discussions and Hussam Abu Safiya for inspiration. RS thanks Salva Duran-Nebreda for early work and useful suggestions on this topic, and the support of a grant AGAUR FI-SDUR 2020, an AEI-PID2023-152129NB-I00 grant, and the Santa Fe Institute. J. P. M. was supported by the PRE2020-091968 grant, funded by MCIN/AEI (10.13039/501100011033), and co-funded by the ESF through the program “Investing in your future”.

