## Supplementary Material for "Animals or plants? Evolutionary branching of sessile versus mobile cognitive agents in noisy environments"

#### I. AGENT-BASED MODEL

The agent-based model presented here is an evolutionary dynamical system in which a set of agents $\{\mathcal{A}_k\}$ exists, perishes, and evolves on a two-dimensional lattice $\Omega$. This lattice is defined as an $N_x \times N_y$ array of discrete sites (patches) that can contain both agents and resources, defined as

$$\Omega = \big\{(i, j) \mid 1 \leq i \leq N_x, 1 \leq j \leq N_y\big\} \in \mathbb{N}^2,$$

with periodic boundary conditions, so that the domain has the topology of a torus and edge effects are eliminated. Each site $(i, j)$ may hold at most one agent at any given time (hard-core exclusion), which implements local competition for space; sites are otherwise unconstrained in the number of resource particles they hold. All distances on the lattice are measured with the $L^1$ (Manhattan) metric, consistent with the four-neighbor structure of the grid; this choice makes both movement and perception follow the natural graph distance of the lattice rather than a straight-line (Euclidean) one.

##### A. Environmental resources

Each patch $(i, j)$ may contain two distinct resources: a non-depletable resource $\mathcal{L}(i, j)$ and a depletable resource $\mathcal{R}(i, j, t)$. The non-depletable resource acts as an analog of "light" from which energy can be harvested and is spatially deterministic, and temporally constant; whereas the particulate resource is spatially heterogeneous, temporally stochastic, and rival (consumption removes particles). Both resources need metabolic investment with an associated cost in order to process them and increase the agent's own mass.

###### Light irradiation

The non-depletable resource $\mathcal{L}(i, j)$ models constant light irradiation, a fixed, time-invariant quantity across the domain $\Omega$. Two spatial distributions for $\mathcal{L}(i, j)$ are examined; the choice of configuration is part of the simulation setup, and all other rules of the model are identical in the two cases.

*Homogeneous configuration.* In the homogeneous configuration, $\mathcal{L}(i, j) = \mathcal{L}$ is constant across the entire domain. This configuration removes spatial variation in the imposed light field and is used to isolate the role of frequency-dependent selection and demographic noise in generating phenotypic differentiation. The light intensity $\mathcal{L}$ is the control parameter that tunes the profitability of autotrophy.

---

*Linear gradient configuration.* In the non-homogeneous configuration, irradiation decreases linearly along one spatial dimension according to the gradient

$$\mathcal{L}(i,j) = \mathcal{L}^{+} - \eta \left| i - \frac{N_x + 1}{2} \right|, \qquad \eta = \frac{2\left(\mathcal{L}^{+} - \mathcal{L}^{-}\right)}{N_x - 1}, \tag{1}$$

where $\left(\mathcal{L}^{+}, \mathcal{L}^{-}\right)$ are the maximum and minimum irradiation levels, and $\eta$ is the gradient slope. With periodic boundary conditions along both axes, the imposed discrete light field has the same minimum value at the lateral seam, $\mathcal{L}(1,j) = \mathcal{L}(N_x, j) = \mathcal{L}^{-}$. It is a V-shaped profile, with highest irradiation $\mathcal{L}^{+}$ at the lattice center and minimum $\mathcal{L}^{-}$ at the lateral edges $i = 1$ and $i = N_x$. Agents crossing the seam pass through the darkest part of the environment. This spatial configuration transforms environmental heterogeneity into a direct driver of geographic niche partitioning: high-irradiation columns select for autotrophic investment, low-irradiation columns for heterotrophic foraging, and the intermediate columns create a transition zone.

#### Depletable resource

The depletable resource $\mathcal{R}(i,j,t) \in \mathbb{N}_0$ signifies the number of discrete particles at site $(i,j)$ at time $t$. Its dynamics are controlled by local, spatially independent, and memoryless stochastic processes for degradation and replenishment. The agent loop is performed asynchronously, followed by the resource update at the end of each time step; the resource update acts independently at every site (agent-mediated depletion is described in Section I.B). The complete update ordering is given in Algorithms 1 and 2.

*Degradation.* The degradation function $\Phi_{\mathrm{d}} : \mathbb{N}_0 \times [0,1] \to \mathbb{N}_0$ captures the resource decay,

$$\Phi_{\mathrm{d}}(r,u) = \begin{cases} 0, & \text{if } u \le \delta_{\mathcal{R}}, \\ r, & \text{otherwise,} \end{cases}$$

effectively reducing to zero the resource $r$ with probability $\delta_{\mathcal{R}}$, in a density-independent manner. This is a "reset" mechanism rather than a linear decay: with probability $\delta_{\mathcal{R}}$ a site is cleared completely, and with probability $1 - \delta_{\mathcal{R}}$ it is left untouched. The reset is memoryless in the sense that the same clearing probability applies every step, so a particle that survived the previous step has no "age" advantage. Physically, this implements catastrophic local depletion that destroys the entire local value at once, rather than a gradual exponential decay.

*Replenishment.* The replenishment function $\Phi_{\mathrm{r}} : [0,1] \to \mathbb{N}_0$ introduces new particles independently of the current resource level. The number of incoming particles at each site is an independent random draw $X$ with mean $\alpha$, the input rate that controls how productive the environment is. The variant used in the main simulations follows a geometric distribution on $\mathbb{N}_0$ with parameter

$$p = \frac{\alpha}{1+\alpha},$$

mean $\alpha$, and variance $\alpha\,(1+\alpha)$. This is generated from a uniform variate $u_{\mathrm{r}}$ via the inverse transform method

$$\Phi_{\mathrm{r}}^{\mathrm{G}}(u) = \left\lfloor \frac{\log(1-u)}{\log(\alpha) - \log(1+\alpha)} \right\rfloor,$$

which satisfies, for all $k \in \mathbb{N}_0$,

$$\mathbb{P}\Big(\Phi_{\mathrm{r}}^{\mathrm{G}}(u_{\mathrm{r}}) = k\Big) = \frac{1}{1+\alpha}\left(\frac{\alpha}{1+\alpha}\right)^{k}.$$

The geometric input has moments

$$\mathbb{E}[X] = \alpha, \qquad \mathrm{Var}(X) = \alpha\,(1+\alpha).$$

Thus $\alpha$ controls both mean input and input variance.

An alternative Bernoulli law, which adds one particle with probability $0 \le \alpha \le 1$ and none otherwise,

$$\Phi_{\mathrm{r}}^{\mathrm{B}}(u) = \begin{cases} 1 & \text{if } u \le \alpha, \\ 0 & \text{otherwise,} \end{cases}$$

has the same mean $\alpha$ but smaller variance $\alpha(1-\alpha)$. It is used as a lower-variance control. Both laws have the same mean input, so differences are purely due to variability: the Bernoulli input has a lighter tail and produces fewer and smaller extreme resource peaks, only due to accumulation over time steps.

It yields the same qualitative phenotypic differentiation as the geometric law but in a narrower light window (see Figure 1). Because sensing is rewarded by extremes, the lighter tail can lower the heterotrophic payoff at fixed $\alpha$; the effect on coexistence boundaries should therefore be assessed from the corresponding simulations.

*Composite update.* The combined update from time $t$ to $t+1$ is given by the composite function

$$\mathcal{R}(i,j,t+1) = \Phi_{\mathrm{r}}(u_{\mathrm{r}}) + \Phi_{\mathrm{d}}\big(\mathcal{R}(i,j,t), u_{\mathrm{d}}\big),$$

where $u_{\mathrm{d}}, u_{\mathrm{r}} \sim \mathcal{U}(0,1)$ are independent random numbers drawn for each site and time step. The reset is applied first and the input second, so that the value of a site is cleared with probability $\delta_{\mathcal{R}}$ before the new particles are added; the stationary statistics derived in Section III correspond to this ordering. This form is used to generate uncertainty in the environment.

In particular, the renewal process is Markovian and spatially independent across sites, and the input is temporally white; because the value persists from one step to the next with probability $1-\delta_{\mathcal{R}}$, the particle field carries a temporal memory on a timescale $\sim \delta_{\mathcal{R}}^{-1}$, while remaining analytically tractable.

*Pre-equilibration.* The agents are introduced into an environment that is already statistically stationary. The resource field is first evolved for a burn-in period of $B$ update steps in the absence of any agent, chosen long enough to reach the stationary state.

### B. Agents

#### Agent state

An agent is characterized by a tuple

$$\mathcal{A}_k = \{m_k, \mathbf{x}_k, \lambda_k, \rho_k\},$$

where $m_k \in \mathbb{R}_0^+$ denotes the agent's mass, and $\mathbf{x}_k \in \Omega$ its spatial location. The variable $\lambda_k \in [0,1]$ represents the fraction of the agent's normalized energy budget allocated to exploiting the non-depletable resource (light); consequently, $1-\lambda_k$ is the fraction directed toward harvesting the depletable resource $\mathcal{R}$. The agent's sensing capability for $\mathcal{R}$ is governed by

$$\rho_k \in \{0, 1, \dots, \rho_{\max}\},$$

which defines an effective search radius $r_k$, defined only for $\rho_k > 0$, $r_k := \rho_k - 1$. If $\rho_k = 0$, the agent is completely oblivious to its environment: it neither perceives nor consumes the depletable resource, and it can only harvest light.

The traits $(\lambda_k, \rho_k)$ are the two evolutionary degrees of freedom of the model. They implement the core trade-off: a large $\lambda$ commits the energy budget to the predictable light resource, which requires no movement or sensing, but forgoes the potentially richer particulate resource; a large $\rho$ grants a wider perceptual neighborhood and hence access to resource-rich patches, but the sensory apparatus costs $c_\rho\, \rho_k$ each step and movement incurs the additional cost $c_d\, d_k$. The allocation parameter $\lambda$ evolves through small Gaussian mutations, whereas $\rho$ evolves through unit steps, reflecting the different resolutions of these traits in the ABM.

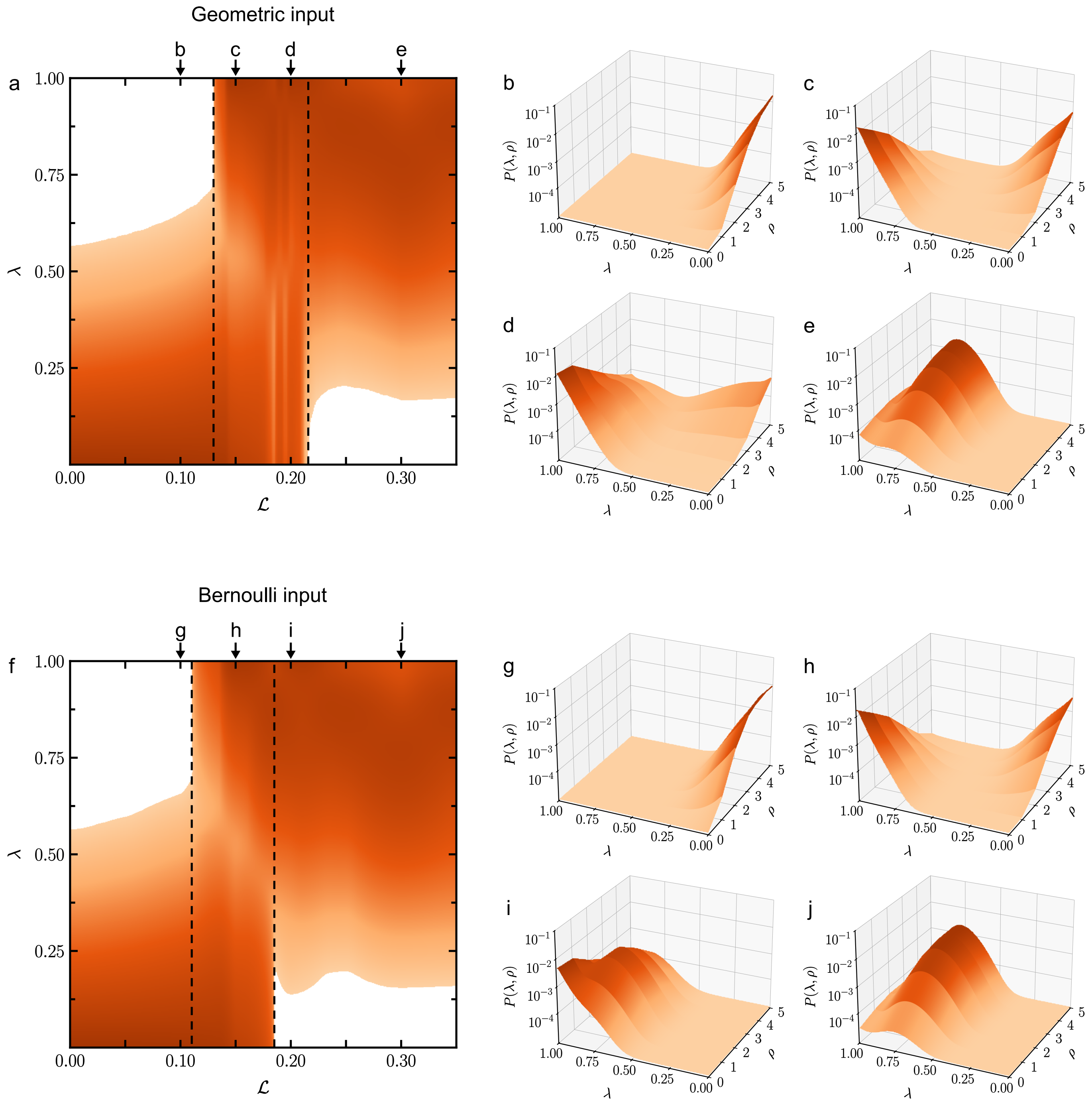


FIG. 1 **Evolution of phenotypic distributions across varying light availability in homogeneous environments: geometric versus Bernoulli resource input.** Panels (a) and (f) display the long-term normalized density distribution of the light-harvesting trait $\lambda$ as a function of uniform environmental light intensity $\mathcal{L}$ for geometric (left column, a–e) and Bernoulli (right column, f–j) input, respectively. The color mapping indicates the marginal probability $P(\lambda)$ on a logarithmic scale, highlighting regions of high phenotypic abundance. Dashed lines delineate the coexistence regime where animal-like and plant-like phenotypes co-occur. Panels (b–e) and (g–j) present cross-sections of the joint probability density $P(\lambda, \rho)$ at selected light levels for geometric and Bernoulli input, respectively, showing the same bimodal structure in both cases with the Bernoulli coexistence window displaced to lower $\mathcal{L}$. All other rules and parameters are identical: $\alpha = 0.5$, $\beta = 1$, $\gamma = 0.1$, $\delta_{\mathcal{A}} = 0.1$, $\delta_{\mathcal{R}} = 0.1$, $\mu = 0.1$, $\rho_{\max} = 5$, $\sigma_\lambda = 0.05$, $c_d = 1$, $c_{\mathcal{L}} = 1$, $c_{\mathcal{R}} = 1$, $c_\rho = 1$.

#### Perception and movement range

Let $\Gamma_k$ denote the set of lattice sites within the perceptual neighborhood

$$\Gamma_k := \big\{(i,j) \mid \|\mathbf{x}_k - (i,j)\|_1 \le r_k\big\},$$

where $\|\cdot\|_1$ is the $L^1$ norm (Manhattan distance). The corresponding set of observable resource patches is

$$\mathcal{R}_k := \big\{\mathcal{R}(i,j,t) \mid (i,j) \in \Gamma_k\big\}.$$

The agent assesses and can reach all patches in $\Gamma_k$, but harvests only at the site it currently occupies. A Manhattan ball of radius $r$ contains $|\Gamma(r)| = 1 + 2r\,(r+1)$ sites; for $\rho \in \{1,2,3,4,5\}$, i.e. $r = \rho - 1 \in \{0,\dots,4\}$, this gives $|\Gamma| \in \{1,5,13,25,41\}$. Thus $\rho = 1$ permits local foraging but no directed movement, which requires $\rho > 1$.

#### Agent dynamics

At each time step, agents are updated asynchronously in a uniformly random order, as detailed in Algorithm 1. Each agent's step is detailed in Algorithm 2.

*Random death.* With probability $\delta_{\mathcal{A}}$, the agent randomly dies. This phenotype-independent mortality provides demographic turnover and space for reproduction. If the agent survives, then resource gathering and potential movement follow.

*Foraging and movement.* For agents with $\rho_k > 1$, acquisition of the depletable resource drives movement; an agent with $\rho_k = 1$ perceives only its own site and therefore remains stationary. Let $\mathcal{X} := \{\mathbf{x}_s \mid \mathcal{A}_s \in \mathcal{A}\}$ denote the set of all agent positions on the lattice. The agent evaluates the legal candidate set

$$N_k := \big(\Gamma_k \setminus \mathcal{X}\big) \cup \{\mathbf{x}_k\},$$

which comprises all unoccupied perceived sites together with the agent's current position (re-included because $\mathbf{x}_k \in \mathcal{X}$). Let $F_k(i,j)$ denote the instantaneous net gain, based on the currently observed resource value, from moving to patch $(i,j)$,

$$F_k(i,j) = \gamma\,(1-\lambda_k)^\beta\,\mathcal{R}(i,j,t) - c_d\,\delta_\mathrm{m}\,\|\mathbf{x}_k - (i,j)\|_1, \tag{2}$$

where $\gamma$ is the resource-to-mass conversion parameter and $c_d\,\delta_\mathrm{m}$ is the cost of moving one lattice unit. The agent chooses a maximizer,

$$(i^*,j^*) = \operatorname*{arg\,max}_{(i,j)\in N_k} F_k(i,j),$$

and moves only when the maximal gain is strictly positive. If several sites tie, the tie is broken uniformly at random; the current site is always a candidate, so an agent never moves merely because all accessible alternatives are non-positive. The resulting movement distance is

$$d_k = \begin{cases} \big\|\mathbf{x}_k - (i^*,j^*)\big\|_1, & \text{if } F_k(i^*,j^*) > 0, \\ 0, & \text{otherwise.} \end{cases}$$

This occupancy rule implements hard-core exclusion during movement: agents cannot displace one another, and spatial competition is resolved through reproduction into empty neighboring sites.

*Mass update.* The agent then updates its mass based on resource intake and metabolic costs. At its current location $(i,j)$, it gains energy from both the non-depletable light resource $\mathcal{L}(i,j)$ and the depletable resource $\mathcal{R}(i,j,t)$. These gains are offset by a baseline maintenance cost and additional costs associated with its allocation strategy, sensing investment, and movement. The net mass change over the time step is given by

$$\Delta m_k = \lambda_k^\beta\,\mathcal{L}(i,j) + \mathbb{1}_{\{\rho_k>0\}}\Big[\gamma\,(1-\lambda_k)^\beta\,\mathcal{R}(i,j,t)\Big] - \delta_\mathrm{m}\,\big[1 + c_{\mathcal{L}}\,\lambda_k + c_{\mathcal{R}}\,(1-\lambda_k) + c_\rho\,\rho_k + c_d\,d_k\big]. \tag{3}$$

The individual terms encode the physiological trade-off explicitly. The light return $\lambda_k^{\beta}\,\mathcal{L}(i,j)$ is deterministic and non-depleting, whereas the heterotrophic return is available only to agents with $\rho_k > 0$, is collected at the current site after movement, and removes the entire local resource value by resetting $\mathcal{R}(i,j,t)$ to zero. The factor $\gamma\,(1-\lambda_k)^{\beta}$ converts the removed resource value into biomass; the remainder, if any, is not retained in the model. The unit baseline term $\delta_{\mathrm{m}}$ is paid by every agent; $c_{\mathcal{L}}\,\lambda_k$ and $c_{\mathcal{R}}\,(1-\lambda_k)$ maintain the two resource-processing capacities; $c_\rho\,\rho_k$ is the continuous sensing cost; and $c_d\,d_k$ is paid only for distance actually traveled. Thus sensing buys information, movement buys access to richer sites, and both investments are favored only when their returns exceed their maintenance costs.

The returns exponent $\beta > 0$ acts on both gross returns and sets the curvature of the energy-allocation trade-off: $\beta = 1$ gives the linear returns used in all agent-based simulations (constant marginal returns per unit of allocation), $0 < \beta < 1$ gives concave returns that favor mixed (intermediate) allocations, and $\beta > 1$ gives convex returns that favor specialists. The Adaptive Dynamics model of Section IV inherits the same exponent and classifies the three regimes.

*Starvation.* If, after the update, the agent's mass becomes non-positive—that is, if $m_k + \Delta m_k \leq 0$—the agent perishes due to starvation. Starvation is the phenotype-dependent death channel in addition to random death.

*Reproduction.* Reproduction is possible only if the agent's mass before clamping meets or exceeds the unit threshold,

$$m_k + \Delta m_k \geq 1.$$

The threshold $m = 1$ defines binary fission. The mass is first clamped to the admissible range

$$[x]_a^b := \min\big(b, \max(a, x)\big),$$

so excess mass above one is not stored. A nearest-neighbor site is then selected uniformly; if it is occupied, reproduction is aborted, whereas an empty site receives the offspring. When reproduction succeeds, the clamped mass is divided equally between parent and offspring, conserving biomass within the lineage. Because the probability of finding an empty neighbor decreases as the lattice fills, this local rule supplies the spatial density dependence (crowding) affecting reproduction rates.

*Mutation.* The offspring inherits the parent's traits $\lambda_k$ and $\rho_k$, which may then undergo mutation. With probability $\mu$, the energy allocation parameter is perturbed by Gaussian noise:

$$\lambda_l = \Big[\lambda_k + \zeta\Big]_0^1, \qquad\qquad \zeta \sim \mathcal{N}(0, \sigma^2).$$

The Gaussian mutation is incremental when $\sigma$ is small relative to the trait range.

Independently, with probability $\mu$, $\rho$ mutates by an equiprobable unit step $\varepsilon \in \{-1, 1\}$, with the result clamped to its admissible range,

$$\rho_l = \Big[\rho_k + \varepsilon\Big]_0^{\rho_{\max}}, \qquad\qquad \varepsilon \in \{-1, 1\}.$$

The Gaussian mutation produces a bounded random walk in $\lambda$, while the unit-step mutation produces a bounded random walk in $\rho$; clamping creates boundary states but is not equivalent to an exactly reflecting random walk.

**Data:** $\Omega$, $\mathcal{L}$, $\mathcal{R}$, $\mathcal{A}$
**Result:** $\mathcal{R}$, $\mathcal{A}$
`// Agent updates (asynchronous)`
Let $K$ be a random permutation of $\{1, \ldots, |\mathcal{A}|\}$
Let $\mathcal{X} \leftarrow \{\mathbf{x}_s \mid \mathcal{A}_s \in \mathcal{A}\}$
**for** $k \in K$ **do**
  `AgentStep(`$k, \mathcal{A}, \mathcal{X}, \mathcal{L}, \mathcal{R}$`)`
**end**
`// Resource update (synchronous)`
**for** $(i, j) \in \Omega$ **do**
  `// Reset (degradation)`
  **if** $\mathcal{U}(0,1) \leq \delta_{\mathcal{R}}$ **then**
    $\mathcal{R}(i,j) \leftarrow 0$
  **end**
  `// Replenishment`
  $u \sim \mathcal{U}(0,1)$
  $\mathcal{R}(i,j) \leftarrow \mathcal{R}(i,j) + \Phi_{\mathrm{r}}(u)$
**end**

**Algorithm 1:** Simulation Step.

**Procedure** `AgentStep(`$k, \mathcal{A}, \mathcal{X}, \mathcal{L}, \mathcal{R}$`):`
  `// Random death`
  **if** $\mathcal{U}(0,1) \leq \delta_{\mathcal{A}}$ **then**
    $\mathcal{A} \leftarrow \mathcal{A} \setminus \{\mathcal{A}_k\}$
    $\mathcal{X} \leftarrow \mathcal{X} \setminus \{\mathbf{x}_k\}$
    **return**
  **end**
  `// Movement`
  $d_k \leftarrow 0$
  **if** $\rho_k > 1$ **then**
    $N_k \leftarrow (\Gamma_k \setminus \mathcal{X}) \cup \{\mathbf{x}_k\}$
    $(i^*, j^*) \leftarrow \arg\max_{(i,j) \in N_k} F_k(i,j)$
    **if** $F_k(i^*, j^*) > 0$ **then**
      $d_k \leftarrow \|\mathbf{x}_k - (i^*, j^*)\|_1$
      $\mathcal{X} \leftarrow (\mathcal{X} \setminus \{\mathbf{x}_k\}) \cup \{(i^*, j^*)\}$
      $\mathbf{x}_k \leftarrow (i^*, j^*)$
    **end**
  **end**
  $(i, j) \leftarrow \mathbf{x}_k$
  `// Mass gain`
  $m_k \leftarrow m_k + \lambda_k^{\beta} \, \mathcal{L}(i,j)$
  **if** $\rho_k > 0$ **then**
    $m_k \leftarrow m_k + \gamma(1 - \lambda_k)^{\beta} \mathcal{R}(i,j)$
    $\mathcal{R}(i,j) \leftarrow 0$
  **end**
  `// Mass loss`
  $C \leftarrow 1 + c_{\mathcal{L}} \lambda_k + c_{\mathcal{R}}(1 - \lambda_k) + c_{\rho} \rho_k + c_d d_k$
  $m_k \leftarrow m_k - \delta_{\mathrm{m}} C$
  `// Starvation death`
  **if** $m_k \leq 0$ **then**
    $\mathcal{A} \leftarrow \mathcal{A} \setminus \{\mathcal{A}_k\}$
    $\mathcal{X} \leftarrow \mathcal{X} \setminus \{\mathbf{x}_k\}$
    **return**
  **end**
  **if** $m_k < 1$ **then**
    **return**
  **end**
  `// Reproduction`
  $m_k \leftarrow \min(m_k, 1)$
  $N \leftarrow \big\{(i,j) \mid \|\mathbf{x}_k - (i,j)\|_1 = 1\big\}$
  $(i', j') \sim \mathcal{U}(N)$
  **if** $(i', j') \in \mathcal{X}$ **then**
    **return**
  **end**
  $m_k \leftarrow m_k / 2$
  $\mathcal{A}_l \leftarrow \mathcal{A}_k$
  $\mathbf{x}_l \leftarrow (i', j')$
  `// Trait mutation`
  **if** $\mathcal{U}(0,1) \leq \mu$ **then**
    $\zeta \sim \mathcal{N}(0, \sigma^2)$
    $\lambda_l \leftarrow \min\big(1, \max(0, \lambda_k + \zeta)\big)$
  **end**
  **if** $\mathcal{U}(0,1) \leq \mu$ **then**
    $\varepsilon \sim \mathcal{U}\big(\{-1, 1\}\big)$
    $\rho_l \leftarrow \min\big(\rho_{\max}, \max(0, \rho_k + \varepsilon)\big)$
  **end**
  $\mathcal{A} \leftarrow \mathcal{A} \cup \{\mathcal{A}_l\}$
  $\mathcal{X} \leftarrow \mathcal{X} \cup \{(i', j')\}$
**end**

**Algorithm 2:** Single-Agent Step.

### II. OBSERVABLES

The observables of this section are evaluated on the ABM trajectories defined above and provide the quantitative criteria used in the main text to identify branching. They also motivate the mean-field quantities approximated in Section III.

#### A. Population classification

To monitor evolutionary divergence, the population is partitioned by the allocation trait at the midpoint of the trait space. Let $\Lambda_- = \{\lambda_k \mid \lambda_k < 1/2\}$ and $\Lambda_+ = \{\lambda_k \mid \lambda_k > 1/2\}$ denote the subsets of allocation traits on either side of the partition, with cardinalities $N_- = |\Lambda_-|$ and $N_+ = |\Lambda_+|$. For a finite set $S = \{x_1, \dots, x_n\}$, let $x_{(1)} \le x_{(2)} \le \cdots \le x_{(n)}$ be the order statistics. The median is defined as

$$\text{median}(S) = \begin{cases} x_{((n+1)/2)} & \text{if } n \text{ is odd}, \\ \frac{1}{2}\left(x_{(n/2)} + x_{(n/2+1)}\right) & \text{if } n \text{ is even}. \end{cases}$$

We define the branch medians as $\tilde{\lambda}_- = \text{median}(\Lambda_-)$ and $\tilde{\lambda}_+ = \text{median}(\Lambda_+)$. The median is used in preference to the arithmetic mean because it minimizes the sum of absolute deviations rather than squared deviations, conferring robustness against transient mutants and outliers that might arise during branching events. The median also remains stable under skewed or multimodal distributions and preserves ordinal consistency with threshold-based classifications.

### III. ANALYTIC APPROXIMATIONS

Because the stochastic rules of Sections I.A–I.B are explicit and Markovian, several ABM quantities can be computed analytically, at least in the mean-field (spatially uncorrelated) approximation. These estimates are not exact for the spatially structured ABM; they provide independent intuition for its viability and sensing behavior. The baseline $\mathcal{R}_0 = \alpha/\delta_{\mathcal{R}}$ is used only within these ABM approximations and is not identified with the resource level of the separate AD model.

#### A. Stationary statistics of the resource field

Consider a single site in the absence of consumption. The renewal is $\mathcal{R}_{t+1} = G_t + Z_t\mathcal{R}_t$, where $G_t$ is the input innovation (independent of the current value) and $Z_t \sim \text{Bernoulli}(1-\delta_{\mathcal{R}})$ indicates survival of the previous value, with $G_t$ and $Z_t$ independent. For any input law with mean $\mathbb{E}[G] = \alpha$, taking expectations gives

$$\mathbb{E}[\mathcal{R}_{t+1}] = \alpha + (1-\delta_{\mathcal{R}})\,\mathbb{E}[\mathcal{R}_t],$$

so the stationary mean—the unexploited resource level in the absence of consumption—is

$$\mathcal{R}_0 := \mathbb{E}[\mathcal{R}^*] = \frac{\alpha}{\delta_{\mathcal{R}}}.$$

The stationary variance depends on the input law through its second moment; the formulas below apply to the specified reset-before-input update ordering. For any input variance $v_G = \text{Var}(G)$, the stationary variance is

$$\text{Var}(\mathcal{R}^*) = \frac{v_G}{\delta_{\mathcal{R}}} + (1-\delta_{\mathcal{R}})\frac{\alpha^2}{\delta_{\mathcal{R}}^2}.$$

For the geometric input, $\text{Var}(G) = \alpha\,(1+\alpha)$, and using the law of total variance—with

$$\text{Var}(Z_t\mathcal{R}_t) = (1-\delta_{\mathcal{R}})\,\text{Var}(\mathcal{R}_t) + (1-\delta_{\mathcal{R}})\,\delta_{\mathcal{R}}\,\mathbb{E}[\mathcal{R}_t]^2$$

—the stationary variance satisfies

$$\text{Var}(\mathcal{R}^*) = \alpha\,(1+\alpha) + (1-\delta_{\mathcal{R}})\,\text{Var}(\mathcal{R}^*) + (1-\delta_{\mathcal{R}})\,\delta_{\mathcal{R}}\,\mathcal{R}_0^2,$$

which yields the closed form

$$\mathrm{Var}(\mathcal{R}^*) = \frac{\alpha\,(1+\alpha)}{\delta_{\mathcal{R}}} + (1-\delta_{\mathcal{R}})\,\frac{\alpha^2}{\delta_{\mathcal{R}}^2}.$$

From these two expressions the relative fluctuation level of the resource field takes the simple form

$$\mathrm{CV} := \frac{\sqrt{\mathrm{Var}(\mathcal{R}^*)}}{\mathbb{E}[\mathcal{R}^*]} = \sqrt{1+\frac{\delta_{\mathcal{R}}}{\alpha}},$$

which depends only on the ratio $\delta_{\mathcal{R}}/\alpha$ and not on the mean resource abundance.

For the Bernoulli input, whose variance $\alpha\,(1-\alpha)$ is smaller, the same computation yields

$$\mathrm{CV}^2 = 1 + \delta_{\mathcal{R}}\left(\frac{1}{\alpha} - 2\right),$$

which is valid for the Bernoulli control with $0 < \alpha \leq 1$ and remains nonnegative; the qualitative conclusions below are therefore insensitive to the choice of input law.

At large input rate $\alpha \gg \delta_{\mathcal{R}}$, $\mathrm{CV} \to 1$, so even the most productive environments retain order-one relative fluctuations—the resource field is never "smooth"; at small input rate $\alpha \ll \delta_{\mathcal{R}}$, $\mathrm{CV} \to \sqrt{\delta_{\mathcal{R}}/\alpha} \gg 1$, so scarce environments are extremely patchy, with occasional large clumps separated by long empty stretches. For the parameter ranges considered, the particle field can be more variable than a Poisson process with the same mean—for which $\mathrm{CV} = 1/\sqrt{\mathcal{R}_0} \ll 1$ at large $\mathcal{R}_0$. This is the sense in which the environment is "noisy": the input law produces order-one relative fluctuations across the parameter ranges considered, so a strategy that relies on the depletable resource must cope with genuine uncertainty, and information about the current field configuration is therefore valuable.

### B. Viability of specialist strategies

Ignoring spatial correlations, competition with other agents, and the movement cost (paid only by moving foragers and added explicitly in the heterotrophic estimate below), the expected per-step mass change of a strategy $(\lambda, \rho)$ is, with linear returns $\beta = 1$,

$$\mathbb{E}[\Delta m] \approx \lambda\mathcal{L} + \mathbb{1}_{\{\rho>0\}}\,\gamma(1-\lambda)\,\mathcal{R}_0 - \delta_{\mathrm{m}}\big[1 + c_{\mathcal{L}}\lambda + c_{\mathcal{R}}(1-\lambda) + c_\rho\rho\big],$$

where the mean local resource value is

$$\mathcal{R}_0 = \frac{\alpha}{\delta_{\mathcal{R}}}.$$

#### Autotrophy

A pure autotroph has $(\lambda, \rho) = (1, 0)$, for which

$$\mathbb{E}[\Delta m] = \mathcal{L} - \delta_{\mathrm{m}}\,(1 + c_{\mathcal{L}}).$$

An isolated autotroph is therefore individually viable whenever the light level exceeds the survival threshold

$$\mathcal{L}_{\mathrm{surv}} := \delta_{\mathrm{m}}\,(1 + c_{\mathcal{L}}).$$

The per-step surplus $\mathcal{L} - \mathcal{L}_{\mathrm{surv}}$ gives the deterministic no-death estimate for the time to reach the reproduction threshold: starting from a sub-threshold mass $m_0 < 1$, an autotroph must gain $1 - m_0$ mass units, requiring approximately

$$T_{\mathrm{aut}} \approx \frac{1 - m_0}{\mathcal{L} - \mathcal{L}_{\mathrm{surv}}}$$

steps when the surplus is positive. Random death, clamping, and failed reproduction are not included in this estimate.

When $\mathcal{L} > \mathcal{L}_{\mathrm{surv}}$, viability alone understates selection near the "plant" phenotype: once a high-$\lambda$ strategy gains enough mass in a single step to reach the fission threshold, further increases in $\lambda$ can no longer raise the division rate. The realized growth rate therefore saturates—even though the per-step surplus keeps growing linearly in $\lambda$—which flattens selection across a range of near-plant phenotypes able to reproduce at every step.

#### Heterotrophy

A forager with $(\lambda, \rho) = (0, \rho)$ and $\rho \geq 1$ harvests $\gamma \mathcal{R}$ from its site and can move toward richer sites. The naive per-step expectation, which ignores the extreme-value gains of movement while still paying the movement cost, is

$$\mathbb{E}[\Delta m] \approx \gamma \mathcal{R}_0 - \delta_{\mathrm{m}} (1 + c_{\mathcal{R}} + c_\rho \rho) - \delta_{\mathrm{m}} c_d \mathbb{E}[d],$$

which is positive under the following mean-field sufficient condition,

$$\frac{\gamma \alpha}{\delta_{\mathcal{R}}} > \delta_{\mathrm{m}} (1 + c_{\mathcal{R}} + c_\rho \rho) + \delta_{\mathrm{m}} c_d \mathbb{E}[d].$$

Heterotrophy is thus viable above a threshold input rate that grows with the sensing trait and with the distance actually traveled; when this threshold is comfortably exceeded, i.e. when $\gamma \mathcal{R}_0$ exceeds the maintenance burden, the forager's per-step net gain is large and its generation time correspondingly short, so heterotrophy is a productive strategy in its own right rather than a marginal one.

This asymmetry—autotrophy is viable whenever light exceeds a fixed threshold, while heterotrophy can be viable when resource input is sufficiently large—helps create conditions under which the two endpoint strategies can be attractors rather than a single optimum.

### C. The benefit of sensing

The key advantage of a larger sensing radius $\rho$ is not captured by the mean-field estimate above, because it derives from the *extreme values* of the particle field rather than its mean.

A forager with radius $r = \rho - 1$ inspects the $|\Gamma(r)| = 1 + 2r(r+1)$ sites of its perceptual neighborhood and selects the one with the largest value. The relevant distribution is that of the stationary field, not of a fresh input draw. Because the renewal process is reset intermittently and accumulates nonnegative inputs, its stationary upper tail is expected to be light-tailed; the following exponential-tail form is used only as a heuristic mean-field approximation,

$$\mathbb{P}(\mathcal{R} > x) \sim \exp\bigl(-x/\mathcal{R}_0\bigr), \qquad \mathcal{R}_0 = \frac{\alpha}{\delta_{\mathcal{R}}},$$

up to factors of order one in the exponent. An exponential tail places the field in the Gumbel domain of attraction, so the maximum over independently drawn sites grows logarithmically in their number:

$$\mathbb{E}\left[\max_{k \leq |\Gamma(r)|} \mathcal{R}_k\right] \approx \mathcal{R}_0 \ln |\Gamma(r)|.$$

(The statement is an order-of-magnitude mean-field estimate: spatial correlations are neglected and the prefactor of the logarithm carries constants of order one.) The benefit of sensing is thus directly proportional to the patchiness of the environment: because the input law yields order-one relative fluctuations (Section III), the gap between the best and the average site is large, and a sensor that samples many sites reaps a large multiple of the mean, which rationalizes why sensing pays precisely in the regime $\alpha \ll \delta_{\mathcal{R}}$ where the field is most variable. This is related to the moving hypothesis [1]: cognitive machinery (sensing) pays off precisely when the environment is uncertain, i.e., when $\alpha$ is small relative to $\delta_{\mathcal{R}}$ and the field is patchy.

Under the independent-site exponential-tail approximation, the same argument explains why the optimal sensing trait is not unbounded: the marginal gain of an extra unit of $\rho$ is

$$\mathcal{R}_0 \ln\left(\frac{|\Gamma(r+1)|}{|\Gamma(r)|}\right),$$

which decreases with $\rho$ (returns to scale in the number of sampled sites are logarithmic), whereas the marginal cost $\delta_{\mathrm{m}} c_\rho$ is constant.

#### D. The branching threshold and the adaptive valley

The heuristic energetics above also rationalize the observation that the light intensity at which plants start "evolving" in the population substantially exceeds the level at which a single autotroph could survive. For an isolated autotroph the viability threshold is $\mathcal{L}_{\text{surv}} = \delta_{\text{m}}\,(1 + c_{\mathcal{L}})$. However, the ancestral population is not composed of autotrophs: it consists of generalists with $\lambda < 0.5$, which (for the case where $\rho = 0$) have a per-step expectation of mass gain

$$\mathbb{E}[\Delta m] = 0.5\,\mathcal{L} - \delta_{\text{m}}\,(1 + 0.5\,c_{\mathcal{L}} + 0.5\,c_{\mathcal{R}}).$$

More generally, the viability condition for an intermediate phenotype $\lambda \in [0,1]$ with $\rho = 0$ is

$$\mathcal{L} > \delta_{\text{m}}\,\frac{1 + c_{\mathcal{R}} + (c_{\mathcal{L}} - c_{\mathcal{R}})\,\lambda}{\lambda},$$

which is a decreasing function of $\lambda$: the higher the allocation to light, the lower the light level at which the strategy is viable.

Denote by $\mathcal{L}_{\text{gen}}$ the generalist's viability threshold, obtained by evaluating the display above at $\lambda = 1/2$:

$$\mathcal{L}_{\text{gen}} = \delta_{\text{m}}(2 + c_{\mathcal{L}} + c_{\mathcal{R}}),$$

which always exceeds the autotrophic threshold by

$$\mathcal{L}_{\text{gen}} - \mathcal{L}_{\text{surv}} = \delta_{\text{m}}\,(1 + c_{\mathcal{R}}) > 0.$$

In this window the autotrophic specialist is viable while the generalist is not. So a plant lineage cannot simply "appear" from nothingness, it must be produced by incremental mutation from the generalist state, and those incremental steps pass through intermediate phenotypes whose viability requires $\mathcal{L}$ well above the survival threshold. At such intermediate $\mathcal{L}$ the generalist must also compete with a heterotrophic population whose members move to the "best" resource sites and thereby capture the surplus energy that would otherwise offset the generalist's more inefficient light harvesting.

The population-level threshold for the appearance of plants may therefore exceed the autotrophic survival condition because the generalist lineage must survive and traverse the intervening trait region while competing with foragers. This is a heuristic interpretation of the ABM mean-field estimates, not an exact threshold.

### IV. ADAPTIVE DYNAMICS MODEL

The trait-based Adaptive Dynamics (AD) model is a separate well-mixed, continuous-trait model [2, 3], analyzed independently of the ABM in the main text (Methods–Adaptive Dynamics). Its parameters are model-specific; the notation is shared for convenience, but numerical values and thresholds need not coincide with those of the ABM. The AD resource equilibrium is defined by its own chemostat equations and is not identified with the ABM approximation $\mathcal{R}_0 = \alpha/\delta_{\mathcal{R}}$. It averages stochastic returns, treats $\rho$ as continuous, replaces explicit spatial uptake and distance costs by a saturating response $\Theta(\rho, \mathcal{R})$ with handling time $\nu$, and represents crowding with logistic regulation. The allocation and sensory traits are analogous to those in the ABM, while the AD model asks whether diversification can arise from physiological trade-offs alone; it is not a mean-field derivation of the ABM.

#### A. Model Formulation

Each individual is characterized by two continuous traits. The allocation fraction $\lambda \in [0,1]$ denotes the proportion of the energy budget devoted to light harvesting (autotrophy); the remaining fraction $1 - \lambda$ is allocated to heterotrophic resource uptake. The foraging radius $\rho \in \mathbb{R}_{\geq 0}$ quantifies the sensing and movement range.

##### Per-capita growth rate

Following the mass update rule of the agent-based model, the instantaneous per-capita growth rate $f$ in the well-mixed AD setting is

$$f(\lambda, \rho; \mathcal{R}) = \mathcal{L}\,\lambda^{\beta} + \gamma\,(1 - \lambda)^{\beta}\,\Theta(\rho, \mathcal{R}) - \delta_{\text{m}}\,\big[1 + c_{\mathcal{L}}\,\lambda + c_{\mathcal{R}}\,(1 - \lambda) + c_{\rho}\,\rho\big]. \tag{4}$$

Here $\mathcal{L}$ is the homogeneous light intensity, $\gamma$ is the resource-to-mass conversion efficiency, and $c_{\mathcal{L}}, c_{\mathcal{R}}, c_\rho$ are maintenance costs for light harvesting, substrate processing, and sensing. The returns exponent $\beta > 0$ acts on both gross returns, whereas the costs remain linear in allocation.

Resource uptake follows a saturating Monod (Holling type II) functional response [4, 5],

$$\Theta(\rho, \mathcal{R}) = \frac{\rho\mathcal{R}}{1 + \nu\,\rho\mathcal{R}}, \tag{5}$$

where the handling time $\nu$ imposes diminishing returns on foraging efficiency. Its first and second derivatives with respect to $\rho$ are

$$\Theta_\rho = \frac{\partial\Theta}{\partial\rho} = \frac{\mathcal{R}}{(1 + \nu\,\rho\mathcal{R})^2}, \qquad \Theta_{\rho\rho} = \frac{\partial^2\Theta}{\partial\rho^2} = -\frac{2\nu\mathcal{R}^2}{(1 + \nu\,\rho\mathcal{R})^3} < 0. \tag{6}$$

##### Effective parameters

Collecting the baseline turnover $\delta_\mathrm{m}$ into the trait-specific costs yields the effective parameters (7):

$$\tilde{c}_{\mathcal{L}} := \delta_\mathrm{m}\, c_{\mathcal{L}}, \qquad \tilde{c}_{\mathcal{R}} := \delta_\mathrm{m}\, c_{\mathcal{R}}, \qquad \tilde{c}_\rho := \delta_\mathrm{m}\, c_\rho, \qquad \delta := \delta_\mathrm{m}, \tag{7}$$

in terms of which the per-capita growth rate becomes (8):

$$f(\lambda, \rho; \mathcal{R}) = \mathcal{L}\,\lambda^\beta + \gamma\,(1-\lambda)^\beta\,\Theta(\rho, \mathcal{R}) - \tilde{c}_{\mathcal{L}}\,\lambda - \tilde{c}_{\mathcal{R}}\,(1-\lambda) - \tilde{c}_\rho\,\rho - \delta. \tag{8}$$

##### Resource dynamics

The depletable resource evolves according to chemostat-like dynamics [6],

$$\frac{\mathrm{d}\mathcal{R}}{\mathrm{d}t} = \alpha - \delta_{\mathcal{R}}\,\mathcal{R} - \iint \Theta(\rho, \mathcal{R})\, n(\lambda, \rho, t)\, \mathrm{d}\rho\, \mathrm{d}\lambda, \tag{9}$$

with $\alpha$ the constant external input rate, $\delta_{\mathcal{R}}$ the combined rate of abiotic decay and dilution, and the integral term capturing the total consumption of the resource by the population across all trait values.

##### Population dynamics

Density dependence is modeled by logistic regulation

$$\frac{\partial n(\lambda, \rho, t)}{\partial t} = \left[ f(\lambda, \rho; \mathcal{R}) \left(1 - \frac{N(t)}{K}\right) - \delta_n \right] n(\lambda, \rho, t), \tag{10}$$

where $N(t) = \iint n(\lambda, \rho, t)\, \mathrm{d}\rho\, \mathrm{d}\lambda$ is the total population density, $K$ is the carrying capacity of the homogeneous environment, and $\delta_n$ denotes the background mortality rate of the density field $n$.

#### B. Ecological equilibrium and invasion fitness

For a monomorphic resident population with phenotype $(\lambda, \rho)$ at ecological equilibrium, the coupled system (9)–(10) reduces to

$$f(\lambda, \rho; \mathcal{R}^*) \left(1 - \frac{N^*}{K}\right) - \delta_n = 0, \tag{11}$$

$$\alpha - \delta_{\mathcal{R}}\,\mathcal{R}^* - \Theta(\rho, \mathcal{R}^*)\, N^* = 0, \tag{12}$$

which jointly determine the equilibrium resource concentration $\mathcal{R}^*$ and population density $N^*$ as functions of the resident phenotype.

The invasion fitness of a rare mutant is its long-run per-capita growth rate in the ecological environment set by the resident population at its own equilibrium. This is the quantity whose sign decides both the direction of the canonical equation and the mutual invasibility of the plant and animal boundaries analyzed thereafter. A rare mutant phenotype $(\lambda_{\mathrm{m}}, \rho_{\mathrm{m}})$ has invasion fitness [2, 3]

$$\sigma(\lambda_{\mathrm{m}}, \rho_{\mathrm{m}}; \lambda, \rho) = f(\lambda_{\mathrm{m}}, \rho_{\mathrm{m}}; \mathcal{R}^*) \left(1 - \frac{N^*}{K}\right) - \delta_n, \tag{13}$$

where $\mathcal{R}^*$ and $N^*$ are the equilibrium resource level and resident population density determined by the resident phenotype $(\lambda, \rho)$. A mutant can invade when $\sigma > 0$; at a resident ecological equilibrium, $\sigma(z; z) = 0$.

#### C. Selection gradient and singular strategy

The direction of evolution is dictated by the selection gradient $\mathbf{g} = (g_\lambda, g_\rho)^{\mathrm{T}}$, the derivative of the invasion fitness with respect to the mutant traits, evaluated at the resident phenotype:

$$g_\lambda(\lambda, \rho) = \left.\frac{\partial \sigma}{\partial \lambda_{\mathrm{m}}}\right|_{\mathrm{m=r}} = \left[\mathcal{L}\,\beta\,\lambda^{\beta-1} - \gamma\,\beta\,(1-\lambda)^{\beta-1}\,\Theta(\rho, \mathcal{R}^*) - \tilde{c}_{\mathcal{L}} + \tilde{c}_{\mathcal{R}}\right] \left(1 - \frac{N^*}{K}\right), \tag{14}$$

$$g_\rho(\lambda, \rho) = \left.\frac{\partial \sigma}{\partial \rho_{\mathrm{m}}}\right|_{\mathrm{m=r}} = \left[\gamma\,(1-\lambda)^{\beta}\,\Theta_\rho(\rho, \mathcal{R}^*) - \tilde{c}_\rho\right] \left(1 - \frac{N^*}{K}\right). \tag{15}$$

The allocation gradient now contains the marginal returns of both resources. The light marginal is $\mathcal{L}\,\beta\,\lambda^{\beta-1}$, while increasing $\lambda$ removes heterotrophic capacity at marginal cost $\gamma\,\beta\,(1-\lambda)^{\beta-1}\,\Theta$. For $\beta > 1$, both returns are convex and the marginal contrast becomes stronger near the two boundaries; for $0 < \beta < 1$, both are concave and intermediate allocations are favored by curvature.

Evolution proceeds through incremental mutational steps according to the canonical equation [3]

$$\frac{\mathrm{d}}{\mathrm{d}t}\begin{pmatrix}\lambda \\ \rho\end{pmatrix} = \mu \begin{pmatrix}\sigma_\lambda^2 & 0 \\ 0 & \sigma_\rho^2\end{pmatrix} \mathbf{g}(\lambda, \rho), \tag{16}$$

where $\mu$ is the mutation rate and $\sigma_\lambda^2$, $\sigma_\rho^2$ denote the mutational variances of the two traits.

A singular strategy $(\lambda^*, \rho^*)$ satisfies $\mathbf{g}(\lambda^*, \rho^*) = \mathbf{0}$, i.e. the simultaneous vanishing of (14) and (15).

##### Stationarity of the allocation gradient

Setting (14) to zero yields

$$\gamma\,\beta\,(1-\lambda^*)^{\beta-1}\,\Theta(\rho^*, \mathcal{R}^*) = \Phi_\beta(\lambda^*), \qquad \Phi_\beta(\lambda) := \mathcal{L}\,\beta\,\lambda^{\beta-1} - \tilde{c}_{\mathcal{L}} + \tilde{c}_{\mathcal{R}}. \tag{17}$$

At an interior singular point the two marginal allocation returns balance. In particular, $\Phi_\beta(\lambda^*) > 0$ is required, and the corresponding uptake level is

$$\Theta^* := \Theta(\rho^*, \mathcal{R}^*) = \frac{\Phi_\beta(\lambda^*)}{\gamma\,\beta\,(1-\lambda^*)^{\beta-1}}.$$

Substituting the Monod functional response (5),

$$\frac{\rho^*\,\mathcal{R}^*}{1 + \nu\,\rho^*\,\mathcal{R}^*} = \Theta^* \qquad \Longrightarrow \qquad \rho^*\,\mathcal{R}^* = \frac{\Theta^*}{1 - \nu\,\Theta^*}, \tag{18}$$

with feasibility requiring $0 < \Theta^* < \nu^{-1}$, or equivalently $0 < \Phi_\beta(\lambda^*) < \gamma\,\beta\,(1-\lambda^*)^{\beta-1}\,\nu^{-1}$.

#### Stationarity of the sensing gradient

Setting (15) to zero gives

$$\gamma\,(1-\lambda^*)^\beta\,\Theta_\rho(\rho^*,\mathcal{R}^*) = \tilde{c}_\rho. \tag{19}$$

Using $\Theta_\rho = \Theta(1-\nu\,\Theta)\,\rho^{-1}$ and the definition of $\Theta^*$, the singular radius is

$$\rho^* = \frac{\gamma\,(1-\lambda^*)^\beta\,\Theta^*\,(1-\nu\,\Theta^*)}{\tilde{c}_\rho} = \frac{\Phi_\beta(\lambda^*)\,(1-\lambda^*)\,(1-\nu\,\Theta^*)}{\beta\,\tilde{c}_\rho}. \tag{20}$$

#### Closing the system

Equations (17)–(20) express the singular heterotrophic gain $\Theta^*$, the product $\rho^*\,\mathcal{R}^*$, and the singular foraging radius $\rho^*$ in terms of $\lambda^*$ and the parameters; the ecological equilibrium (11)–(12) evaluated at the singular point provides the closing condition.

The resident fitness at the singular point is

$$f^* = \mathcal{L}\,(\lambda^*)^\beta + \gamma\,(1-\lambda^*)^\beta\,\Theta^* - \tilde{c}_\mathcal{L}\,\lambda^* - \tilde{c}_\mathcal{R}\,(1-\lambda^*) - \tilde{c}_\rho\,\rho^* - \delta. \tag{21}$$

Using the equivalence $\gamma\,\beta\,(1-\lambda^*)^{\beta-1}\,\Theta^* = \Phi_\beta(\lambda^*)$, this can also be written as

$$f^* = \mathcal{L}\,(\lambda^*)^\beta + \frac{1-\lambda^*}{\beta}\,\Phi_\beta(\lambda^*) - \tilde{c}_\mathcal{L}\,\lambda^* - \tilde{c}_\mathcal{R}\,(1-\lambda^*) - \tilde{c}_\rho\,\rho^* - \delta. \tag{22}$$

The ecological equilibrium and the steady state population concentration $N^*$ then close the system numerically,

$$N^* = \frac{\alpha - \delta_\mathcal{R}\,\mathcal{R}^*}{\Theta^*}.$$

The next subsections classify the evolutionary behavior of this AD model. The mutant-trait Hessian characterizes disruptive versus stabilizing curvature, the resident-gradient Jacobian adds eco-evolutionary feedback through $\mathcal{R}^*$, and the subsequent boundary analysis examines the plant and animal optima. These conclusions apply to the AD model and are not direct predictions of the ABM.

### D. Evolutionary branching analysis

#### Hessian matrix

The Hessian of the invasion fitness (13) with respect to the mutant traits, evaluated at the singular point, is

$$H = \begin{pmatrix} h_{\lambda\lambda} & h_{\lambda\rho} \\ h_{\rho\lambda} & h_{\rho\rho} \end{pmatrix} = \left.\begin{pmatrix} \dfrac{\partial^2\sigma}{\partial\lambda_\mathrm{m}^2} & \dfrac{\partial^2\sigma}{\partial\lambda_\mathrm{m}\partial\rho_\mathrm{m}} \\ \dfrac{\partial^2\sigma}{\partial\lambda_\mathrm{m}\partial\rho_\mathrm{m}} & \dfrac{\partial^2\sigma}{\partial\rho_\mathrm{m}^2} \end{pmatrix}\right|_{(\lambda^*,\rho^*)}, \tag{23}$$

with entries

$$h_{\lambda\lambda} = \left[\mathcal{L}\,\beta\,(\beta-1)\,(\lambda^*)^{\beta-2} + \gamma\,\beta\,(\beta-1)\,(1-\lambda^*)^{\beta-2}\,\Theta(\rho^*,\mathcal{R}^*)\right]\left(1-\frac{N^*}{K}\right), \tag{24}$$

$$h_{\rho\rho} = \gamma\,(1-\lambda^*)^\beta\,\Theta_{\rho\rho}(\rho^*,\mathcal{R}^*)\left(1-\frac{N^*}{K}\right) < 0, \tag{25}$$

$$h_{\lambda\rho} = -\gamma\,\beta\,(1-\lambda^*)^{\beta-1}\,\Theta_\rho(\rho^*,\mathcal{R}^*)\left(1-\frac{N^*}{K}\right) < 0. \tag{26}$$

Both gross returns contribute to the allocation curvature. Writing $Q^* := 1 - N^* K^{-1} > 0$ (which, by the resident balance (11), equals $\delta_n/f^*$, so every curvature entry scales with this small ratio and only signs and relative magnitudes matter),

$$h_{\lambda\lambda} = Q^* \beta (\beta - 1) \left[ \mathcal{L} (\lambda^*)^{\beta-2} + \gamma (1 - \lambda^*)^{\beta-2} \Theta^* \right],$$

whose sign is controlled by $\beta - 1$. The cross term is also modified by the same exponent,

$$h_{\lambda\rho} = -Q^* \gamma \beta (1 - \lambda^*)^{\beta-1} \Theta^*_\rho.$$

The mutant-trait Hessian can therefore be written as

$$H = \left(1 - \frac{N^*}{K}\right) \begin{pmatrix} \mathcal{L} \beta (\beta - 1) (\lambda^*)^{\beta-2} + \gamma \beta (\beta - 1) (1 - \lambda^*)^{\beta-2} \Theta^* & -\gamma \beta (1 - \lambda^*)^{\beta-1} \Theta^*_\rho \\ -\gamma \beta (1 - \lambda^*)^{\beta-1} \Theta^*_\rho & \gamma (1 - \lambda^*)^{\beta} \Theta^*_{\rho\rho} \end{pmatrix}. \tag{27}$$

#### Evolutionary stability

The eigenvalues $\kappa$ of the Hessian (23) satisfy the characteristic equation

$$\kappa^2 - (h_{\lambda\lambda} + h_{\rho\rho}) \kappa + (h_{\lambda\lambda} h_{\rho\rho} - h^2_{\lambda\rho}) = 0. \tag{28}$$

The singular point is an evolutionarily stable strategy (ESS)—$H$ is negative definite—if and only if both eigenvalues are negative. For a symmetric $2 \times 2$ matrix this requires [7, 8]: (i) $h_{\lambda\lambda} < 0$ and $h_{\rho\rho} < 0$; and (ii)

$$\det H = h_{\lambda\lambda} h_{\rho\rho} - h^2_{\lambda\rho} > 0.$$

Since $h_{\rho\rho} < 0$ always, condition (i) requires $h_{\lambda\lambda} < 0$, i.e. $\beta < 1$.

For $\beta > 1$, both allocation contributions to $h_{\lambda\lambda}$ are positive in the interior, so $h_{\lambda\lambda} > 0$, while $h_{\rho\rho} < 0$; hence $H$ cannot be negative definite and an evolutionarily stable interior singularity is not possible.

For $\beta = 1$, both gross returns are linear, so $h_{\lambda\lambda} = 0$ but $h_{\lambda\rho} \neq 0$ whenever sensing has a marginal benefit; therefore $\det H < 0$ and the singularity is a saddle.

For $0 < \beta < 1$, $h_{\lambda\lambda} < 0$ and $h_{\rho\rho} < 0$; the singularity is an ESS when $\det H > 0$, and it is a branching point only when $\det H < 0$. This case requires an interior point and positive $Q^*$; boundary singularities must be treated separately.

#### Convergence stability

For a singular point that is convergence stable but not evolutionarily stable, disruptive selection drives evolutionary branching [9]. Convergence stability is a property of the *resident* gradient dynamics rather than of the mutant-fitness curvature alone: it is decided by the Jacobian of the selection gradient

$$g(z) = \nabla_{z_\mathrm{m}} \sigma(z_\mathrm{m}; z)\big|_{z_\mathrm{m}=z}$$

with respect to the resident traits at the singular point, which includes the response of the ecological equilibrium to the resident phenotype.

At a singular point, because $g_i = Q^* f_i$ with $Q^* = 1 - N^* K^{-1}$ and $f_i \equiv \partial f / \partial z_i$, the Jacobian $J = \mathrm{D}g(z^*)$ splits into a direct part, which is exactly the Hessian entries (24)–(26), and an indirect, eco-evolutionary part acting through the equilibrium resource level,

$$J_{ij} = \left.\frac{\partial g_i}{\partial z_j}\right|_{z^*} = \underbrace{Q^* f_{ij}}_{=h_{ij}} + Q^* f_{i\mathcal{R}} \frac{\partial \mathcal{R}^*}{\partial z_j},$$

where $h_{ij}$ are the Hessian entries (24)–(26), and

$$f_{ij} = \partial^2 f / \partial z_i \partial z_j, \qquad f_{i\mathcal{R}} = \partial^2 f / \partial z_i \partial \mathcal{R}.$$

Two simplifications occur at the singular point. First,

$$\frac{\partial g_i}{\partial N^*} = -\frac{f_i}{K} = 0,$$

because the selection gradient vanishes ($f_i = 0$). Second, the equilibrium resource level is first-order insensitive to the allocation trait: implicitly differentiating the ecological equilibrium (11)–(12) with respect to the resident trait $z_j$ gives the linear system

$$f_j Q^* - \frac{f}{K}\frac{\partial N^*}{\partial z_j} + f_{\mathcal{R}} Q^* \frac{\partial \mathcal{R}^*}{\partial z_j} = 0, \qquad -(\delta_{\mathcal{R}} + \Theta_{\mathcal{R}} N^*)\frac{\partial \mathcal{R}^*}{\partial z_j} - \Theta \frac{\partial N^*}{\partial z_j} = \Theta_\rho \, N^* \, \mathbb{1}_{\{j=\rho\}}, \tag{29}$$

where $\mathbb{1}_{\{j=\rho\}}$ is the indicator function (equal to one when $j = \rho$, zero otherwise), $\Theta_{\mathcal{R}} = \rho^* \, (1 + \nu \, \rho^* \, \mathcal{R}^*)^{-2}$ is the resource derivative of the functional response, and $f_{\mathcal{R}} = \gamma \, (1 - \lambda^*)^\beta \, \Theta_{\mathcal{R}} > 0$ is the marginal resource value of uptake.

At the singular point $f_\lambda = f_\rho = 0$, and provided the ecological equilibrium is regular with $f > 0$, Eq. (29) gives

$$\frac{\partial \mathcal{R}^*}{\partial \lambda} = \frac{\partial N^*}{\partial \lambda} = 0, \qquad \frac{\partial \mathcal{R}^*}{\partial \rho} = -\frac{\Theta_\rho N^*}{\Delta}, \qquad \frac{\partial N^*}{\partial \rho} = \frac{f_{\mathcal{R}} Q^* K}{f} \frac{\partial \mathcal{R}^*}{\partial \rho}, \tag{30}$$

with

$$\Delta = \delta_{\mathcal{R}} + \Theta_{\mathcal{R}} \, N^* + \Theta \, f_{\mathcal{R}} \, Q^* \, \frac{K}{f} > 0.$$

At the singular point, under these regularity and feasibility assumptions, the resource equilibrium responds only to the sensory trait, and only through depletion,

$$\frac{\partial \mathcal{R}^*}{\partial \rho} = -\frac{\Theta_\rho N^*}{\Delta} < 0.$$

while the allocation trait feeds back exclusively through the cross channel $J_{\lambda\rho}$.

Substituting (30) into the Jacobian gives the closed form

$$J = \begin{pmatrix} h_{\lambda\lambda} & h_{\lambda\rho} + c_\lambda \\ h_{\lambda\rho} & h_{\rho\rho} + c_\rho \end{pmatrix}, \tag{31}$$

with the feedback corrections

$$c_\lambda = \frac{Q^* \, \gamma \, \beta \, (1 - \lambda^*)^{\beta - 1} \, \Theta_{\mathcal{R}} \, \Theta_\rho \, N^*}{\Delta} > 0, \qquad c_\rho = -\frac{Q^* \, \gamma \, (1 - \lambda^*)^\beta \, \Theta_{\rho\mathcal{R}} \, \Theta_\rho \, N^*}{\Delta}, \tag{32}$$

where

$$\Theta_{\rho\mathcal{R}} = \frac{1 - \nu \, \rho^* \, \mathcal{R}^*}{(1 + \nu \, \rho^* \, \mathcal{R}^*)^3}$$

is the mixed curvature of the Monod response. The corrections affect only the $\rho$-column, in agreement with (30), and make the Jacobian asymmetric, $J_{\lambda\rho} \neq J_{\rho\lambda}$. Because $\Delta > \Theta_{\mathcal{R}} N^*$ when $f_{\mathcal{R}} > 0$, the coupling correction never reverses the sign of the cross entry,

$$J_{\lambda\rho} = h_{\lambda\rho} + c_\lambda = -Q^* \, \gamma \, \beta \, (1 - \lambda^*)^{\beta-1} \, \Theta_\rho \left(1 - \frac{\Theta_{\mathcal{R}} \, N^*}{\Delta}\right) < 0,$$

whereas the self-correction $c_\rho$ changes sign with $\Theta_{\rho\mathcal{R}}$: using the singular relation

$$\nu \, \rho^* \, \mathcal{R}^* = \frac{\nu \, \Theta^*}{1 - \nu \, \Theta^*},$$

the feedback strengthens the (negative) sensory curvature below the inflection point of the functional response, $\Theta^* < (2 \, \nu)^{-1}$, and weakens it above, $\Theta^* > (2\nu)^{-1}$.

Convergence stability is decided by the linearized canonical equation (16), $\mathrm{d}\xi/\mathrm{d}t = MJ\,\xi$ for a small deviation $\xi$ of the resident trait, with $M = \mu\,\mathrm{diag}(\sigma_\lambda^2, \sigma_\rho^2)$: the singular point is convergence stable if and only if $MJ$ is Hurwitz, i.e. for the $2 \times 2$ system,

$$\det J > 0, \qquad \mu\Big(\sigma_\lambda^2 J_{\lambda\lambda} + \sigma_\rho^2 J_{\rho\rho}\Big) < 0, \tag{33}$$

which for isotropic mutation variance ($\sigma_\lambda = \sigma_\rho$) reduces to $\operatorname{tr} J < 0$ and $\det J > 0$.

The three regimes of $\beta$ then follow. For $\beta > 1$, $J_{\lambda\lambda} = h_{\lambda\lambda} > 0$. If $\det J \le 0$, condition (33) fails; if $\det J > 0$, the negativity of both cross entries forces $J_{\rho\rho} > 0$, and the weighted trace in (33) is positive. In either case the interior singular point is *not* convergence stable for $\beta > 1$: the allocation direction repels, so the singularity cannot be a classical branching point. For $\beta = 1$, $h_{\lambda\lambda} = 0$ gives $J_{\lambda\lambda} = 0$, and since both cross entries are strictly negative,

$$\det J = -J_{\lambda\rho}\,J_{\rho\lambda} < 0,$$

so, whenever the interior point exists with positive cross-derivatives, it is a saddle of the canonical equation and is not convergence stable. The linear-return case therefore realizes branching as *escape from an interior saddle*: the monomorphic population is driven off the singularity along the trade-off axis—the unstable eigenvector has

$$\frac{v_\lambda}{v_\rho} = \frac{J_{\lambda\rho}}{\kappa_+} < 0,$$

the same plant–animal direction identified from the Hessian—and coexistence is then maintained by the mutual invasibility of the two boundary specialists established below. The split does not need a converge-first phase. Two facts produce it. First, the indefinite Hessian makes selection around the generalist disruptive: mutants sufficiently close to the singular point on either side of the disruptive eigenvector have positive invasion fitness (the quadratic form dominated by the positive eigenvalue $\kappa_+$), so a population with any phenotypic spread is pulled apart along the plant–animal axis. Second, the two boundary specialists are mutually invasible, so once either side is reached the other can re-invade and the pair is protected. The split is thus disruptive selection near a repelling saddle, together with protected coexistence of the two boundary optima; in the stochastic ABM, phenotypic spread is supplied by mutation and finite population size. A spatially explicit extension can produce analogous branches, but their coexistence must be established from that model's own dynamics. For $0 < \beta < 1$, $h_{\lambda\lambda} < 0$ and $h_{\rho\rho} < 0$, but convergence stability still requires the additional conditions in Eq. (33). For arbitrary positive mutational variances, the weighted-trace condition in (33) requires $J_{\lambda\lambda} < 0$ and $J_{\rho\rho} < 0$ separately: the first holds automatically here, because $\partial\mathcal{R}^*/\partial\lambda = 0$ at the singular point gives $J_{\lambda\lambda} = h_{\lambda\lambda} < 0$, while the second holds whenever the feedback correction $c_\rho$ does not overwhelm the direct sensory saturation $h_{\rho\rho}$. Combined with the Hessian classification above, the $0 < \beta < 1$ regime is the only one in which the interior point can be a genuine convergence-stable branching point (when $\det H < 0$) or an ESS (when $\det H > 0$).

#### Summary of branching conditions

The "branching" conditions for the three regimes of $\beta$ are summarized as follows. For $\beta > 1$, the interior singularity is not an ESS because the allocation curvature is positive, and the convergence-stability analysis of Section IV.D shows that it is never convergence stable (the allocation direction repels), so it cannot act as a classical branching point. For $\beta = 1$, the cross-trait term makes the Hessian indefinite at an interior singular point whenever $\Theta_\rho^* > 0$, and the full resident-gradient Jacobian has $\det J < 0$: the interior point is a saddle of the canonical equation, so the linear-return regime's branching mechanism is realized as escape from an interior saddle along the plant–animal axis rather than as convergence-stable branching. For $0 < \beta < 1$, branching is conditional: it requires

$$h_{\lambda\rho}^2 > h_{\lambda\lambda}\,h_{\rho\rho},$$

where all three entries now contain the factors $(1-\lambda^*)^{\beta-1}$ or $(1-\lambda^*)^\beta$ shown above.

Thus, convex returns ($\beta > 1$) penalize intermediate allocations and promote specialization, while concave returns ($0 < \beta < 1$) favor mixed allocations unless the allocation–sensing coupling is sufficiently strong. The linear case $\beta = 1$ has no direct allocation curvature, but the cross derivative remains and produces the saddle point that can lead to a branching event.

Intuitively, the concave-returns regime $0 < \beta < 1$ is the only one in which the *converge* phase of classical branching can exist. Convergence requires the selection gradient to point inward, which requires diminishing returns in every trait direction: the marginal benefit of further investment in a channel must decline with investment. This is exactly what Eq. (24) shows for the allocation trait when $\beta < 1$ ($h_{\lambda\lambda} < 0$, because $\beta - 1 < 0$ while both terms in the bracket are positive), and the Monod response delivers for the sensory radius in every regime ($h_{\rho\rho} < 0$). With both self-curvatures negative, the interior mixotroph is a local fitness maximum in each single-trait direction, but this alone does not establish convergence stability; the Jacobian conditions in Eq. (33) must also hold. In the linear-return regime $\beta = 1$ the allocation curvature vanishes ($h_{\lambda\lambda} = 0$): the gradient is flat in $\lambda$, the interior is a saddle ($\det J = -J_{\lambda\rho}\, J_{\rho\lambda} < 0$), and no convergence phase exists. At $\beta > 1$ the allocation curvature is positive ($h_{\lambda\lambda} > 0$): increasing returns make the interior allocation direction locally repelling, although global trajectories and boundary outcomes require separate analysis. Concave returns thus create the attractive generalist on which classical branching depends; whether the converged generalist is then split rather than being an ESS is decided by the coupling condition $h_{\lambda\rho}^2 > h_{\lambda\lambda}\, h_{\rho\rho}$ of the Hessian classification above.

#### Classical-branching window in the concave-returns regime

A finite classical-branching window, if present, must satisfy both defining conditions of a Geritz branching point—convergence stability and evolutionary instability—within the concave-returns regime. The qualitative classification of an interior mixotroph follows from the local conditions derived above: (i) for sufficiently strong diminishing returns, it can be a convergence-stable ESS when the stabilizing self-curvatures dominate the allocation–sensing coupling, $\det H > 0$; (ii) for an intermediate concave-return regime, it can be a genuine branching point when it is convergence stable ($\det J > 0$ with negative weighted trace) but not evolutionarily stable ($\det H < 0$); (iii) as the interior point approaches a feasibility boundary, the divergence between plant and animal is instead described by the two-optima mechanism below. The relevant parameter window is bounded by feasibility, convergence stability, and evolutionary-stability conditions.

*Lower edge: the ESS boundary.* The generalist fails to be evolutionarily stable whenever the allocation–sensing coupling overcomes the stabilizing self-curvatures, $h_{\lambda\rho}^2 > h_{\lambda\lambda}\, h_{\rho\rho}$. As $\beta$ increases toward 1, the allocation curvature

$$h_{\lambda\lambda} = Q^*\beta(\beta - 1)[\mathcal{L}(\lambda^*)^{\beta-2} + \gamma(1 - \lambda^*)^{\beta-2}\Theta^*] \quad \text{[Eq. (24)]}$$

shrinks toward zero—the linear-return limit, in which the marginal benefit of further allocation is flat—while the cross term $h_{\lambda\rho}$ remains finite, so the coupling eventually overcomes the self-curvatures and

$$\det H = h_{\lambda\lambda}\, h_{\rho\rho} - h_{\lambda\rho}^2$$

changes sign: any convergence-stable generalist sufficiently close to $\beta = 1$ is necessarily disruptive. Away from the linear-return value the diminishing-returns curvature stabilizes the generalist again, and it is a convergence-stable ESS only where the self-curvatures dominate the coupling. Because this lower edge is a purely local curvature balance at the singular point, it is set by the trade-off geometry rather than by the ecology.

*Upper edge: the feasibility boundary.* As $\beta$ increases toward 1, the singular point slides toward the plant corner of trait space (the heterotrophic allocation and the sensing investment both shrink, $\lambda^* \to 1$, $\rho^* \to 0$), while the equilibrium resource level it requires rises; at some exponent below $\beta = 1$ the interior point collides with the plant vertex and ceases to exist. Above that exponent—and in particular at the linear-return value $\beta = 1$ adopted by the agent-based model—the population cannot converge to any generalist, which is why the linear-return regime realizes the two-optima mechanism rather than classical branching. Unlike the lower edge, the upper edge is an ecological feasibility condition and shifts with the resource supply.

#### Direction of disruptive selection

When the interior Hessian is indefinite, the eigenvector associated with its positive eigenvalue identifies the disruptive direction. For the linear-return case ($\beta = 1$) and, more generally, whenever the cross term dominates with negative sign, this direction points along the plant–animal axis:

$$\frac{v_\lambda}{v_\rho} = \frac{h_{\lambda\rho}}{\kappa_+ - h_{\lambda\lambda}}. \tag{34}$$

For $\beta = 1$, $h_{\lambda\lambda} = 0$ and

$$\kappa_+ = \frac{1}{2}\left(h_{\rho\rho} + \sqrt{h_{\rho\rho}^2 + 4\,h_{\lambda\rho}^2}\right) > 0,$$

so

$$\frac{v_\lambda}{v_\rho} = \frac{h_{\lambda\rho}}{\kappa_+} < 0,$$

high allocation is paired with low sensing, and vice versa. For other exponents, the same interpretation applies when the positive Hessian eigenvector has opposite-sign components; it should not be asserted solely from the sign of $\beta - 1$.

#### Two notions of branching

Two distinct phenomena are conventionally both called branching, and only one is meant by the classical term: (i) *Classical (Geritz) branching* [3, 9], an interior singular point of the canonical equation that is convergence stable but not evolutionarily stable—the population first converges to the generalist and is then split by disruptive selection into two diverging branches that eventually reach the opposite boundaries of trait space. This requires a genuine convergence phase: the singular point must be an attractor of the canonical equation, $\det J > 0$ and $\operatorname{tr} J < 0$ under isotropic mutation; (ii) *Two-optima branching*, the fixed-environment fitness landscape has two local maxima—here the plant boundary $(\lambda, \rho) = (1, 0)$ and the animal boundary $(\lambda, \rho) = (0, \rho_a)$—and the interior balance point between them is a repelling saddle of the canonical equation. A monomorphic population started anywhere near the interior is pushed away from the generalist, so no convergence phase exists; if the two optima are additionally mutually invasible through the ecology each creates, both can be realized and persist as a protected dimorphism.

The model can realize the second mechanism at $\beta = 1$ and may do so at $\beta > 1$, subject to boundary feasibility and mutual invasibility. In the linear-return regime ($\beta = 1$), an interior point with the stated nonzero cross-derivatives is a saddle, $\det J = -J_{\lambda\rho}\, J_{\rho\lambda} < 0$; for $\beta > 1$ the allocation direction repels, $J_{\lambda\lambda} > 0$. In neither case is the point convergence stable, so no classical branching point exists—yet the model does branch, as escape from the interior along the trade-off axis toward the two boundary optima, and the mutual invasibility established below makes the coexistence protected. A spatially explicit extension can produce the same two branches; once produced, their coexistence is mediated by movement and local competition rather than by an explicitly modeled diffusion process.

Only in the concave-returns regime $0 < \beta < 1$ can the interior point be convergence stable under the stated interior assumptions; there the model can exhibit classical (Geritz) branching when the Hessian is indefinite ($h_{\lambda\rho}^2 > h_{\lambda\lambda}\, h_{\rho\rho}$).

### E. The two specialist steady states

The Hessian and Jacobian analysis establishes when the interior generalist is disruptive but does not specify the attractors that replace it. The analysis above identifies conditions for local diversification but leaves open where trajectories eventually go. The candidate evolutionary endpoints are a plant-like branch with $(\lambda, \rho) \to (1, 0)$ and an animal-like branch with $\lambda \to 0$ and $\rho > 0$. Their boundary accessibility and ecological feasibility are analyzed below: the plant allocation is a boundary value, while the animal radius $\rho_a$ is an interior root of the $\rho$-gradient when such a root exists. Their endpoint fitnesses are independent of $\beta$, because $\lambda^\beta = 1$ at $\lambda = 1$ and $(1 - \lambda)^\beta = 1$ at $\lambda = 0$; however, their boundary stability conditions do depend on the one-sided derivatives, which can vanish, remain finite, or diverge as $\beta$ changes. The closed forms below therefore describe the endpoint ecological states, while their evolutionary accessibility must be checked using the appropriate one-sided gradient, and the next subsection tests whether those endpoints can coexist.

#### Plant-like steady state

A plant harvests no particulate resource ($\Theta(0,\mathcal{R})=0$), so the equilibrium resource level is the unexploited level

$$\mathcal{R}_p^* = \mathcal{R}_0 \equiv \frac{\alpha}{\delta_{\mathcal{R}}},$$

and the per-capita growth rate at the steady state is

$$f_p = f(1,0;\mathcal{R}_p^*) = \mathcal{L} - \tilde{c}_{\mathcal{L}} - \delta,$$

with no dependence on $\beta$. The population steady state follows from the logistic factor $(1-N/K)$ of the AD population dynamics [Eq. (10)],

$$N_p^* = K\left(1 - \frac{\delta_n}{f_p}\right),$$

and is positive whenever $f_p > \delta_n$, i.e.

$$\mathcal{L} > \tilde{c}_{\mathcal{L}} + \delta + \delta_n.$$

With $\delta = \delta_{\mathrm{m}}$ and $\tilde{c}_{\mathcal{L}} = \delta_{\mathrm{m}}\, c_{\mathcal{L}}$ this reproduces the ABM survival threshold $\mathcal{L}_{\mathrm{surv}} = \delta_{\mathrm{m}}\,(1 + c_{\mathcal{L}})$ of Section III, augmented by the background mortality $\delta_n \approx \delta_{\mathcal{A}}$. At the plant boundary, the resident has $\rho = 0$, so $\Theta(0,\mathcal{R}_0) = 0$, and the fixed-$\rho = 0$ one-sided allocation gradient is

$$g_{\lambda,p}^{-} = Q_p\,[\mathcal{L}\,\beta - \tilde{c}_{\mathcal{L}} + \tilde{c}_{\mathcal{R}}], \qquad Q_p = 1 - \frac{N_p^*}{K}.$$

The sensing gradient is $g_{\rho,p} = -Q_p\,\tilde{c}_\rho < 0$ at $\lambda = 1$, so the plant boundary is locally stable against increasing sensing, provided the boundary interpretation $\rho \geq 0$ is used. Attraction toward the allocation boundary additionally requires $g_{\lambda,p}^{-} > 0$ under the convention that positive selection increases $\lambda$. A mutant that changes both $\lambda$ and $\rho$ follows a joint boundary direction and must be assessed from the full invasion fitness rather than by inserting an unspecified $\rho$ into the allocation derivative. Thus endpoint attraction is a one-sided condition and cannot be inferred from endpoint fitness alone. When the plant boundary is attracting, its ecological state is the one given above.

#### Animal-like steady state

At $\lambda = 0$ the light term vanishes for every $\beta > 0$, and the sensory trait relaxes to the value at which the $\rho$-gradient vanishes, $\partial\sigma/\partial\rho_{\mathrm{m}} = 0$:

$$\gamma\,\Theta_\rho(\rho_a,\mathcal{R}_a^*) = \tilde{c}_\rho, \qquad \Theta_\rho(\rho,\mathcal{R}) = \frac{\mathcal{R}}{(1+\nu\,\rho\,\mathcal{R})^2}.$$

Solving for the radius gives the closed form

$$\rho_a = \frac{\sqrt{\gamma\,\mathcal{R}_a^*/\tilde{c}_\rho} - 1}{\nu\,\mathcal{R}_a^*},$$

which is positive whenever the equilibrium value satisfies $\gamma\,\mathcal{R}_a^* > \tilde{c}_\rho$; otherwise the optimum is the boundary value $\rho_a = 0$. The corresponding heterotrophic gain is

$$\Theta_a = \Theta(\rho_a,\mathcal{R}_a^*) = \frac{1 - \sqrt{\tilde{c}_\rho/(\gamma\,\mathcal{R}_a^*)}}{\nu},$$

and the per-capita growth rate is $f_a = \gamma\,\Theta_a - \tilde{c}_{\mathcal{R}} - \tilde{c}_\rho\,\rho_a - \delta$. The ecological equilibrium closes the system: the resource balance $\alpha - \delta_{\mathcal{R}}\mathcal{R}^* - \Theta_a N^* = 0$ fixes the animal population

$$N_a^* = \frac{\alpha - \delta_{\mathcal{R}}\,\mathcal{R}_a^*}{\Theta_a},$$

and the logistic balance $f_a(1 - N_a^*/K) = \delta_n$ reduces to a single equation for the equilibrium resource level,

$$\left[\gamma\,\Theta_a - \tilde{c}_{\mathcal{R}} - \tilde{c}_\rho\,\rho_a - \delta\right]\left[1 - \frac{\alpha - \delta_{\mathcal{R}}\,\mathcal{R}_a^*}{K\,\Theta_a}\right] = \delta_n,$$

with $\rho_a$ and $\Theta_a$ given by the closed forms above; this equation is transcendental in $\mathcal{R}_a^*$ (through the square roots), and a feasible steady state additionally requires $N_a^* > 0$, i.e. $\alpha > \delta_{\mathcal{R}}\,\mathcal{R}_a^*$, and $f_a > \delta_n$. In the mean-field approximation of Section III, which evaluates the resource at its unexploited level $\mathcal{R}_0 = \alpha/\delta_{\mathcal{R}}$, these reduce to the fully explicit estimates

$$\rho_a \simeq \frac{\sqrt{\gamma\,\mathcal{R}_0/\tilde{c}_\rho} - 1}{\nu\,\mathcal{R}_0}, \qquad \Theta_a \simeq \frac{1 - \sqrt{\tilde{c}_\rho/(\gamma\,\mathcal{R}_0)}}{\nu}.$$

The animal boundary is attracting when the one-sided allocation gradient for mutants entering from above is negative. At $\lambda = 0$,

$$g^{+}_{\lambda,a} = Q_a\left[\mathcal{L}\,\beta\,\lambda^{\beta-1} - \gamma\,\beta\,(1-\lambda)^{\beta-1}\,\Theta_a - \tilde{c}_{\mathcal{L}} + \tilde{c}_{\mathcal{R}}\right]_{\lambda\downarrow 0}.$$

For $\beta > 1$, this reduces to $Q_a\,(-\gamma\,\beta\,\Theta_a - \tilde{c}_{\mathcal{L}} + \tilde{c}_{\mathcal{R}})$ when the one-sided derivative is evaluated at $\lambda = 0^+$; for $\beta = 1$, attraction requires $\gamma\,\Theta_a > \mathcal{L} + \tilde{c}_{\mathcal{R}} - \tilde{c}_{\mathcal{L}}$; and for $0 < \beta < 1$, the positive light marginal diverges, so the animal boundary is not attracting under the unconstrained continuous-trait model. In all cases the viability condition remains $f_a > \delta_n$.

#### F. Mutual invasibility of the two steady states

The preceding one-strain analysis is now used to test the frequency-dependent coexistence of the two boundaries. When the mutual-invasibility inequalities below hold, each specialist can invade the ecological equilibrium of the other. This is the local protected-coexistence criterion for the two-specialist AD system and represents negative frequency-dependent selection rather than spatial structure in the ABM. The criterion complements the local invasibility result above and identifies the parameter window relevant to the coexistence regions of the agent-based phase diagram. The invasion fitness of a rare mutant into a resident population at its ecological equilibrium is [Eq. (13)]

$$\sigma(\lambda_{\mathrm{m}}, \rho_{\mathrm{m}}; \lambda, \rho) = f(\lambda_{\mathrm{m}}, \rho_{\mathrm{m}}; \mathcal{R}^*)\left(1 - \frac{N^*}{K}\right) - \delta_n,$$

and a mutant invades whenever $\sigma > 0$.

##### The animal invades the plant steady state

With the plant as resident, the particulate resource stands at the unexploited level $\mathcal{R}_p^* = \mathcal{R}_0 \equiv \alpha/\delta_{\mathcal{R}}$, and the plant's logistic balance gives

$$1 - \frac{N_p^*}{K} = \frac{\delta_n}{f_p}.$$

An animal-like mutant carrying the trait $\rho_a$ of the animal steady state has invasion fitness

$$\sigma_{a\to p} = f(0, \rho_a; \mathcal{R}_0)\left(1 - \frac{N_p^*}{K}\right) - \delta_n = \delta_n\left(\frac{f(0, \rho_a; \mathcal{R}_0)}{f_p} - 1\right),$$

and therefore invades whenever $f(0, \rho_a; \mathcal{R}_0) > f_p$, i.e.

$$\gamma\,\Theta(\rho_a, \mathcal{R}_0) - \tilde{c}_{\mathcal{R}} - \tilde{c}_\rho\,\rho_a > \mathcal{L} - \tilde{c}_{\mathcal{L}}.$$

Because the plant does not consume $\mathcal{R}$, the animal mutant faces the largest possible value $\mathcal{R}_0$, so this is the least restrictive condition the heterotrophic strategy can encounter: since $\Theta$ increases with $\mathcal{R}$, $f(0, \rho; \mathcal{R})$ is increasing in $\mathcal{R}$ for any fixed $\rho$.

#### The plant invades the animal steady state

With the animal as resident, the resource is depleted to $\mathcal{R}_a^*$ and the logistic balance gives

$$1-\frac{N_a^*}{K}=\frac{\delta_n}{f_a}.$$

The plant's growth rate does not depend on the resource level, $f(1,0;\mathcal{R}_a^*)=\mathcal{L}-\tilde{c}_{\mathcal{L}}-\delta=f_p$, so

$$\sigma_{p\to a}=f_p\left(1-\frac{N_a^*}{K}\right)-\delta_n=\delta_n\left(\frac{f_p}{f_a}-1\right),$$

and the plant invades whenever $f_p>f_a$, i.e.

$$\mathcal{L}-\tilde{c}_{\mathcal{L}}>\gamma\,\Theta_a-\tilde{c}_{\mathcal{R}}-\tilde{c}_\rho\,\rho_a.$$

In the animal's depleted environment the heterotrophic gain is reduced while the sensing upkeep $\tilde{c}_\rho\,\rho_a$ is still paid; this is precisely the gap that leaves room for the autotrophic strategy.

#### The coexistence window

Subject to the feasibility and boundary-attraction conditions above, the two steady states are mutually invasive whenever the net autotrophic return lies strictly between the animal's net heterotrophic gain evaluated in its own (depleted) environment and in the plant's unexploited environment:

$$\gamma\,\Theta_a-\tilde{c}_{\mathcal{R}}-\tilde{c}_\rho\,\rho_a<\mathcal{L}-\tilde{c}_{\mathcal{L}}<\gamma\,\Theta(\rho_a,\mathcal{R}_0)-\tilde{c}_{\mathcal{R}}-\tilde{c}_\rho\,\rho_a.$$

The window is non-empty in the coexistence regime, because consumption depresses the resource below the unexploited level, $\mathcal{R}_a^*<\mathcal{R}_0$, and $\Theta$ is strictly increasing in $\mathcal{R}$, so the upper bound exceeds the lower bound by the depletion advantage $\gamma\,[\Theta(\rho_a,\mathcal{R}_0)-\Theta_a]$; the window closes only when the animal's depletion of the resource is negligible. The lower inequality is compatible with the animal-boundary dominance condition—$\gamma\,\Theta_a-\tilde{c}_{\mathcal{R}}>\mathcal{L}-\tilde{c}_{\mathcal{L}}$, i.e. the animal would out-earn the plant in its own depleted environment if sensing were free—precisely when the sensing upkeep exceeds the animal's gross net advantage over the plant,

$$\tilde{c}_\rho\,\rho_a>\gamma\,\Theta_a-\tilde{c}_{\mathcal{R}}-(\mathcal{L}-\tilde{c}_{\mathcal{L}})>0.$$

The coexistence window is therefore the regime in which cognition is costly enough that neither specialist can exclude the other. The lower inequality is the condition that the plant's net return beats the animal's net gain in the depleted environment, while the upper inequality expresses that, against an autotrophic resident, the animal finds a resource-rich environment in which sensing pays. Both specialists are therefore best responses to the environment created by the other—the signature of negative frequency-dependent selection and protected coexistence. At the endpoint traits, the endpoint fitness values are independent of $\beta$; the one-sided accessibility conditions are not. The mutual-invasibility inequalities themselves therefore apply directly to the linear-return case $\beta=1$, but their use for other $\beta$ requires the corresponding boundary-accessibility conditions.

### G. Protected coexistence under mutual invasibility

The pairwise invasibility inequalities of the previous subsection are the ecological prerequisite for the dimorphism. For the two-specialist extension considered here, the feasibility and stability analysis below determines when these inequalities yield a positive locally stable coexistence equilibrium.

Consider the ecological dynamics of the two specialists, with densities $p$ (plant-like, trait $(1,0)$) and $a$ (animal-like, trait $(0,\rho_a)$), total density $N=p+a$, and shared resource $\mathcal{R}$,

$$\begin{aligned}\frac{\mathrm{d}p}{\mathrm{d}t}&=\left[f_p\left(1-\frac{N}{K}\right)-\delta_n\right]p,\\ \frac{\mathrm{d}a}{\mathrm{d}t}&=\left[f_a(\mathcal{R})\left(1-\frac{N}{K}\right)-\delta_n\right]a,\\ \frac{\mathrm{d}\mathcal{R}}{\mathrm{d}t}&=\alpha-\delta_{\mathcal{R}}\mathcal{R}-\Theta(\rho_a,\mathcal{R})\,a,\end{aligned}\tag{35}$$

where

$$f_p = \mathcal{L} - \tilde{c}_{\mathcal{L}} - \delta, \qquad f_a(\mathcal{R}) = \gamma\,\Theta(\rho_a, \mathcal{R}) - \tilde{c}_{\mathcal{R}} - \tilde{c}_\rho\,\rho_a - \delta,$$

so the animal's growth rate increases with the resource level, while the plant's is independent of it,

$$\frac{\mathrm{d}f_p}{\mathrm{d}\mathcal{R}} = 0, \qquad \frac{\mathrm{d}f_a}{\mathrm{d}\mathcal{R}} = \gamma\,\frac{\partial\Theta}{\partial\mathcal{R}} > 0,$$

##### Existence

At an interior equilibrium, the per-capita growth rates of both species must be zero. Assuming $f_p > \delta_n$, the logistic slack is positive, and the resource level at the animal–plant coexistence equilibrium must satisfy

$$f_a(\mathcal{R}^*_{\mathrm{ap}}) = f_p,$$

using $\cdot_{\mathrm{ap}}$ to denote animal–plant coexistence.

The total consumer density at this equilibrium is given by

$$N^*_{\mathrm{ap}} = K\left(1 - \frac{\delta_n}{f_p}\right).$$

Because $f_a$ is strictly increasing in $\mathcal{R}$ whenever $\rho_a > 0$, and because the mutual-invasibility conditions imply

$$f_a(\mathcal{R}^*_a) = f_a < f_p < f_a(\mathcal{R}_0),$$

where $f_a \equiv f_a(\mathcal{R}^*_a)$ is the animal's growth rate at its own single-species steady state, the coexistence resource level is uniquely determined as

$$\mathcal{R}^*_{\mathrm{ap}} = f_a^{-1}(f_p) \in (\mathcal{R}^*_a, \mathcal{R}_0).$$

The corresponding animal density is obtained from the resource balance equation:

$$a^*_{\mathrm{ap}} = \frac{\alpha - \delta_{\mathcal{R}}\,\mathcal{R}^*_{\mathrm{ap}}}{\Theta(\rho_a, \mathcal{R}^*_{\mathrm{ap}})} > 0.$$

To verify that the plant density $p^*_{\mathrm{ap}} = N^*_{\mathrm{ap}} - a^*_{\mathrm{ap}}$ is positive, we examine the function

$$h(\mathcal{R}) = \frac{\alpha - \delta_{\mathcal{R}}\,\mathcal{R}}{\Theta(\rho_a, \mathcal{R})}.$$

For $\mathcal{R} < \mathcal{R}_0$, this function is strictly decreasing, since

$$h'(\mathcal{R}) = \frac{-\delta_{\mathcal{R}}\,\Theta - (\alpha - \delta_{\mathcal{R}}\,\mathcal{R})\,\partial\Theta/\partial\mathcal{R}}{\Theta^2} < 0.$$

Consequently,

$$a^*_{\mathrm{ap}} = h(\mathcal{R}^*_{\mathrm{ap}}) < h(\mathcal{R}^*_a) = N^*_a < N^*_{\mathrm{ap}},$$

where the last inequality follows from $f_p > f_a$. Hence, $p^*_{\mathrm{ap}} > 0$. Under the stated inequalities and feasibility conditions, a unique interior equilibrium with both species present therefore exists.

#### Stability

To examine local stability, we linearize the system around the coexistence equilibrium $(p^*_{\rm ap}, a^*_{\rm ap}, \mathcal{R}^*_{\rm ap})$. Linearization involves approximating the nonlinear dynamics by a first-order Taylor expansion near the equilibrium; the resulting Jacobian matrix determines whether small perturbations decay (stability) or grow (instability) over time.

At the equilibrium, we define several positive composite quantities that capture the strength of key ecological interactions. These are evaluated at the coexistence point and will appear as entries in the Jacobian:

$$\bar{p} = \frac{f_p\, p^*_{\rm ap}}{K}, \qquad \bar{a} = \frac{f_p\, a^*_{\rm ap}}{K}.$$

Here, $\bar{p}$ and $\bar{a}$ represent the per-capita density-dependent mortality or self-limitation terms for the plant and animal, respectively. They arise from the logistic growth term and scale with each species' equilibrium density relative to the common carrying capacity $K$. These terms quantify how much each species' growth is reduced by total consumer crowding.

Let us define some quantities to simplify the subsequent analysis.

Let $m$ measure the positive feedback from the resource to the animal's growth rate,

$$m := F'(\mathcal{R}^*_{\rm ap}) \left(1 - \frac{N^*_{\rm ap}}{K}\right) a^*_{\rm ap}$$

The term $F'(\mathcal{R}) > 0$ is the slope of the animal's optimized growth function with respect to resource availability; it captures how efficiently the animal converts additional resource into population growth.

Let $t$ be the animal's resource uptake or consumption rate at equilibrium,

$$t := \Theta(\rho_a, \mathcal{R}^*_{\rm ap})$$

A larger $t$ means stronger consumptive pressure on the resource by the animal population.

Let $s$ represent the total effective loss rate of the resource,

$$s := \delta_{\mathcal{R}} + \left.\frac{\partial \Theta}{\partial \mathcal{R}}\right|_{\mathcal{R}^*_{\rm ap}} a^*_{\rm ap}$$

This combines background resource loss with the additional depletion caused by higher resource levels stimulating greater consumption.

All these quantities are positive under the model assumptions, which ensures that the subsequent stability analysis yields meaningful results.

The Jacobian matrix of the linearized system takes the form

$$J = \begin{pmatrix} -\bar{p} & -\bar{p} & 0 \\ -\bar{a} & -\bar{a} & m \\ 0 & -t & -s \end{pmatrix}.$$

Its characteristic polynomial is

$$\det(\lambda I - J) = \lambda^3 + A\lambda^2 + B\lambda + C,$$

with positive coefficients

$$A = \bar{p} + \bar{a} + s, \qquad B = \bar{p}\,s + \bar{a}\,s + m\,t, \qquad C = \bar{p}\,m\,t.$$

The Routh–Hurwitz criterion for cubic polynomials requires $A > 0$, $C > 0$, and $AB - C > 0$. We compute

$$A\,B - C = (\bar{p} + \bar{a} + s)\,(\bar{p}\,s + \bar{a}\,s + m\,t) - \bar{p}\,m\,t.$$

Expanding and simplifying,

$$A\,B - C = (\bar{p} + \bar{a} + s)\,(\bar{p}\,s + \bar{a}\,s) + m\,t\,(\bar{a} + s) > 0,$$

since all quantities involved are positive. Thus, all Routh–Hurwitz conditions are satisfied, and the coexistence equilibrium is locally asymptotically stable.

#### H. Invasion implies fixation for nearby strategies

The protected coexistence established requires the two specialists to be well separated in trait space. Locally, and under the competitive-exclusion assumptions of this well-mixed model, two sufficiently close phenotypes generically have opposite first-order invasion fitnesses: one is favored and the other is eliminated if the resident is replaced. This local result is the Adaptive Dynamics statement often called invasion implies fixation [3, 9].

Let $z = (\lambda, \rho)$ denote the resident phenotype and $z' = z + \xi$ a nearby mutant with small $\|\xi\|$. Because a resident has zero per-capita growth rate at its own ecological equilibrium, the invasion fitness satisfies

$$\sigma(z; z) = 0 \qquad \text{for all } z,$$

so expanding $\sigma(z'; z)$ and its reverse $\sigma(z; z')$ around the diagonal $z' = z$ yields

$$\sigma(z'; z) = g(z) \cdot \xi + \mathcal{O}(\|\xi\|^2), \qquad \sigma(z; z') = -g(z) \cdot \xi + \mathcal{O}(\|\xi\|^2),$$

where $g(z) = \nabla_{z_\mathrm{m}} \sigma(z_\mathrm{m}; z)|_{z_\mathrm{m}=z}$ is the selection gradient and the dot denotes the scalar product in trait space. The antisymmetry of the linear terms is not an assumption: differentiating the identity $\sigma(z; z) \equiv 0$ along the diagonal gives $\partial\sigma/\partial z_\mathrm{m} = -\partial\sigma/\partial z_\mathrm{r}$ at $z_\mathrm{m} = z_\mathrm{r} = z$, so the coefficient of $\xi$ in $\sigma(z; z')$ is exactly the negative of that in $\sigma(z'; z)$. (The indirect effects through the resident's equilibrium $\mathcal{R}^*$, $N^*$ enter through the resident derivative of $\sigma$ in the second expansion, but the identity $\sigma(z; z) \equiv 0$ pins that derivative to $-g(z)$ on the diagonal, so the first-order antisymmetry is exact rather than approximate.)

Whenever the gradient does not vanish, $g(z) \neq 0$, and the mutation step is sufficiently small that the linear term dominates the quadratic remainder—along a fixed direction $\hat{u} = \xi/\|\xi\|$ with $g(z)\cdot\hat{u} \neq 0$, for $\|\xi\| < 2\,|g(z)\cdot\hat{u}|/|\hat{u}^\mathrm{T} H(z)\hat{u}|$, with $H(z)$ the Hessian of the invasion fitness evaluated at the resident point—the two invasion fitnesses have *opposite* signs:

$$g(z) \cdot \xi > 0 \implies \sigma(z'; z) > 0 \quad \text{and} \quad \sigma(z; z') < 0,$$

and conversely for $g(z) \cdot \xi < 0$. If the mutant invades and the ecological dynamics remain in the local competitive-exclusion regime, the resident cannot reinvade the mutant's monomorphic environment: the mutant grows from rarity while the resident's density declines. If instead the mutant cannot invade, it is eliminated when rare. Thus, for a generic sufficiently small displacement away from a non-singular resident, the first-order selection gradient determines which phenotype is locally favored; protected polymorphism requires a finite trait separation or a singular point where the linear term vanishes. (A displacement exactly perpendicular to the gradient, $g(z) \cdot \xi = 0$, is the degenerate measure-zero case in which both invasion fitnesses are of second order and the outcome is decided by the Hessian; it is never realized by the Gaussian mutation kernel of the ABM.)

At a singular point, $g(z^*) = 0$, both invasion fitnesses are of order $\|\xi\|^2$. The linear dominance above fails, and mutual invasibility can become possible for mutants on opposite sides of the singularity; whether it does so is determined by the Hessian and the ecological feedback. This is the onset of disruptive selection and branching characterized by the Hessian classification above, and it is why the two specialists introduced below can become mutually invasive only after they have diverged to opposite boundaries of trait space.

### V. THREE TRAITS UNDER A SHARED BUDGET

In the Adaptive Dynamics model of Section IV the two allocation traits share a normalized budget, $\lambda$ for light and $1 - \lambda$ for the depletable resource with $\lambda + (1 - \lambda) = 1$, while the sensing radius $\rho$ is paid for separately through the linear cost $\tilde{c}_\rho\, \rho$ and does not compete for that budget. Here we relax this asymmetry and place all three investments—photosynthetic machinery, heterotrophic machinery and sensing—on a single shared budget $B$ weighted by the same cost coefficients. Under the conditions derived below, this formulation can reproduce the same two-optima structure and protected coexistence; the result is not independent of the budget normalization for every parameter choice.

#### Model and budget rescaling

Each individual is now characterized by three non-negative traits—the photosynthetic investment $\lambda_p$, the heterotrophic investment $\lambda_a$, and the foraging radius $\rho$—constrained by the budget

$$\tilde{c}_\mathcal{L}\, \lambda_p + \tilde{c}_\mathcal{R}\, \lambda_a + \tilde{c}_\rho\, \rho \leq B, \tag{36}$$

where $\tilde{c}_{\mathcal{L}}, \tilde{c}_{\mathcal{R}}, \tilde{c}_\rho$ are the effective costs of (7). The per-capita growth rate generalizes (8) to

$$f(\lambda_p, \lambda_a, \rho; \mathcal{R}) = \mathcal{L}\,\lambda_p^\beta + \gamma\,\lambda_a^\beta\,\Theta(\rho, \mathcal{R}) - \tilde{c}_{\mathcal{L}}\,\lambda_p - \tilde{c}_{\mathcal{R}}\,\lambda_a - \tilde{c}_\rho\,\rho - \delta, \tag{37}$$

with the same functional response (5) and the resource and population dynamics (9)–(10). Rescaling each trait by its own cost,

$$\omega_p := \tilde{c}_{\mathcal{L}}\,\lambda_p, \qquad \omega_a := \tilde{c}_{\mathcal{R}}\,\lambda_a, \qquad \omega_\rho := \tilde{c}_\rho\,\rho,$$

turns (36) into the simplex $\omega_p + \omega_a + \omega_\rho \le B$ and (37) into

$$f(\omega_p, \omega_a, \omega_\rho; \mathcal{R}) = L\,\omega_p^\beta + G\,\omega_a^\beta\,\Theta(\omega_\rho, \mathcal{R}) - \omega_p - \omega_a - \omega_\rho - \delta, \tag{38}$$

with

$$L := \frac{\mathcal{L}}{\tilde{c}_{\mathcal{L}}^\beta}, \qquad G := \frac{\gamma}{\tilde{c}_{\mathcal{R}}^\beta\,\tilde{c}_\rho}, \qquad \kappa := \frac{\nu}{\tilde{c}_\rho}, \qquad \Theta(\omega_\rho, \mathcal{R}) = \frac{\omega_\rho\,\mathcal{R}}{1 + \kappa\,\omega_\rho\,\mathcal{R}}. \tag{39}$$

Every unit of budget then costs one unit of fitness. All trait asymmetries are absorbed into the rescaled return coefficients $L$, $G$ and handling time $\kappa$, and the external light level $\mathcal{L}$ enters only through $L$. At an optimum the budget binds, $\omega_p + \omega_a + \omega_\rho = B$, whenever at least one compartment has marginal return above one at that optimum; otherwise the optimum lies strictly inside the simplex and part of the budget is left unspent. Apart from the degenerate zero optimum at $\omega = (0, 0, 0)$ when no compartment pays for itself, such slack with positive investment can only occur in the concave case discussed below.

##### Linear returns: the heterotrophic compartment has increasing returns

At $\beta = 1$ the plant term is linear and the heterotrophic term couples the two heterotrophic traits,

$$f = (L-1)\,\omega_p + \big[G\,\omega_a\,\Theta(\omega_\rho, \mathcal{R}) - \omega_a - \omega_\rho\big] - \delta. \tag{40}$$

The bracket is the only coupling: harvest is proportional to the product $\omega_a\Theta$, so sensing $\omega_\rho$ and machinery $\omega_a$ are complements. For a fixed heterotrophic budget $b = \omega_a + \omega_\rho$ let

$$h(b) := \max_{\omega_a + \omega_\rho = b} \big[G\,\omega_a\,\Theta(\omega_\rho, \mathcal{R})\big] - b.$$

The maximizer and value are in closed form:

$$w = \frac{\sqrt{1 + \kappa\,b\,\mathcal{R}} - 1}{\kappa}, \qquad \omega_\rho^\star = \frac{w}{\mathcal{R}}, \qquad \omega_a^\star = b - \frac{w}{\mathcal{R}}, \qquad h(b) = \frac{G\,w^2}{\mathcal{R}} - b, \tag{41}$$

and, with $y := \sqrt{1 + \kappa\,b\,\mathcal{R}}$,

$$h'(b) = \frac{G\,(y-1)}{\kappa\,y} - 1, \qquad h''(b) = \frac{G\,\mathcal{R}}{2\,y^3} > 0. \tag{42}$$

Thus $h$ is strictly convex over the feasible range, i.e. it has increasing returns, which reflects the complementarity between machinery and sensing. Writing $b = B - \omega_p$ for the budget left to heterotrophy after allocating $\omega_p$ to the plant,

$$\phi(\omega_p) := (L-1)\,\omega_p + h(B - \omega_p)$$

is convex on $[0, B]$ for the stated parameter range and therefore attains its maximum at an endpoint. Under this condition, the optimal phenotype is a single specialist,

$$(\omega_p^\star, \omega_a^\star, \omega_\rho^\star) = \begin{cases} (B, 0, 0), & \text{pure plant}, \quad \mathcal{R} \le \mathcal{R}_{\text{sw}}(B), \\ \Big(0, B - \frac{w}{\mathcal{R}}, \frac{w}{\mathcal{R}}\Big), & \text{optimal heterotroph}, \quad \mathcal{R} \ge \mathcal{R}_{\text{sw}}(B), \end{cases} \tag{43}$$

with $w = \left(\sqrt{1+\kappa B \mathcal{R}} - 1\right)/\kappa$. The interior point that would equalize the three marginal returns,

$$\frac{\partial f}{\partial \omega_p} = \frac{\partial f}{\partial \omega_a} = \frac{\partial f}{\partial \omega_\rho},$$

is the minimum of $\phi$, an anti-optimum, for the unconstrained linear-return reduction.

The pure plant has $\Theta(0, \mathcal{R}) = 0$, hence

$$f_p(B) = L B^\beta - B - \delta,$$

independent of $\mathcal{R}$.

The optimally partitioned heterotroph has

$$f_a^{\text{opt}}(\mathcal{R}) = \frac{G w^2}{\mathcal{R}} - B - \delta,$$

which is nondecreasing in $\mathcal{R}$, and is strictly increasing whenever the optimized heterotrophic allocation is nonzero. The $\mathcal{R}$ at which the two specialists are equally fit is

$$\mathcal{R}_{\text{sw}}(B) = \frac{y^2 - 1}{\kappa B}, \qquad y = \frac{G + \kappa L}{G - \kappa L}, \tag{44}$$

which requires $G > \kappa L$. If $G \leq \kappa L$ the heterotrophic return never compensates its cost and the plant is optimal at every $\mathcal{R}$ in a fixed environment. When $G > \kappa L$, the plant is optimal for $\mathcal{R} < \mathcal{R}_{\text{sw}}(B)$ and the optimally partitioned heterotroph for $\mathcal{R} > \mathcal{R}_{\text{sw}}(B)$. Thus, in a fixed environment and under the convexity condition stated above, the reduced objective has endpoint optima (the plant corner and the optimized heterotroph edge) separated by an interior minimum when that minimum exists. This yields a two-optima structure in this shared-budget model when both endpoint optima are locally viable.

#### Two-optima branching and protected coexistence

The resource level is set by the resident phenotype. The plant does not consume $\mathcal{R}$ and leaves the unexploited level $\mathcal{R}_p = \alpha/\delta_{\mathcal{R}}$; the optimally partitioned heterotroph depletes $\mathcal{R}$ to $\mathcal{R}_a(B) < \mathcal{R}_p$, the root of (9) at $(\omega_a^\star, \omega_\rho^\star)$. Comparing these feedback levels with (44) gives three regimes:

1. If $\mathcal{R}_p \leq \mathcal{R}_{\text{sw}}(B)$ the plant is optimal even in the richest environment it creates and the monomorphic plant is realized,
2. If $\mathcal{R}_a(B) \geq \mathcal{R}_{\text{sw}}(B)$ the heterotroph is optimal even when it has depleted the resource and the monomorphic heterotroph is realized,
3. If $\mathcal{R}_a(B) < \mathcal{R}_{\text{sw}}(B) < \mathcal{R}_p$, no single phenotype is optimal.

In the third regime each specialist is fitter in the environment created by the other—in the plant's world $\mathcal{R}_p > \mathcal{R}_{\text{sw}}$ the heterotroph can invade, and in the heterotroph's world $\mathcal{R}_a < \mathcal{R}_{\text{sw}}$ the plant can invade—so the two optima are mutually invasible and the population is held in protected coexistence by negative frequency dependence, within this shared-budget model.

Assuming that all biological parameters are nonnegative, and restricting attention to feasible equilibria with positive densities of both specialists, the candidate coexistence equilibrium is determined as follows. The resource level $\mathcal{R}_{\text{ap}}^*$ uniquely solves the condition

$$f_a^{\text{opt}}(\mathcal{R}_{\text{ap}}^*) = f_p,$$

where $f_p = LB - B - \delta$ evaluated at $\beta = 1$ (i.e., when the plant invests fully in its own growth). The strict monotonicity of $f_a^{\text{opt}}$ ensures that this solution, when it exists, is unique. The corresponding total consumer density at equilibrium is

$$N_{\text{ap}}^* = K\left(1 - \frac{\delta_n}{f_p}\right),$$

which is positive provided $f_p > \delta_n$. This total density is then partitioned between the animal and plant species according to the resource balance:

$$a^*_{\rm ap} = \frac{\alpha - \delta_{\mathcal{R}}\,\mathcal{R}^*_{\rm ap}}{\Theta(\omega^\star_\rho, \mathcal{R}^*_{\rm ap})}, \qquad p^*_{\rm ap} = N^*_{\rm ap} - a^*_{\rm ap} > 0,$$

where the positivity of $p^*_{\rm ap}$ is guaranteed under the feasibility conditions established earlier.

For a strictly positive coexistence equilibrium with an interior optimized heterotrophic allocation—meaning that the animal invests positive amounts in both sensing and uptake machinery—we have $\Theta > 0$, $\Theta_{\mathcal{R}} > 0$ (the resource uptake function increases with resource availability), and $F'(\mathcal{R}^*) > 0$ (the animal's optimized growth rate responds positively to resource increases). Under these conditions, the local asymptotic stability of the equilibrium follows directly from the corresponding $(p, a, \mathcal{R})$ Jacobian matrix and the Routh–Hurwitz conditions derived in the stability analysis.

Thus, the shared-budget model is capable of reproducing the two-optima structure, provided the above conditions are met. The essential ingredient enabling this outcome is the complementarity between the animal's sensing ability and its heterotrophic machinery: these two components jointly generate increasing returns to the heterotrophic budget, meaning that additional investment in one enhances the returns from the other. This positive synergy creates the nonlinearities necessary for the coexistence of two distinct specialists.

##### Nonlinear returns

Examine the curvature properties of the allocation objective

$$\Psi(\omega_p, \omega_a, \omega_\rho; \mathcal{R}) = L\omega_p^\beta + G\omega_a^\beta \Theta(\omega_\rho, \mathcal{R}),$$

which determines whether interior solutions (generalists) or boundary solutions (specialists) are optimal. The allocation problem involves three trait components: $\omega_p$, $\omega_a$, and $\omega_\rho$, subject to the budget constraint $\omega_p + \omega_a + \omega_\rho \leq B$.

For concave returns ($0 < \beta < 1$), each isolated allocation term exhibits diminishing marginal returns. The plant term $L\,\omega_p^\beta$ is concave in $\omega_p$, and, holding $\omega_a$ fixed, the heterotrophic return is concave in $\omega_\rho$ since $\Theta_{\rho\rho} < 0$. However, the coupled heterotrophic term $G\,\omega_a^\beta\,\Theta(\omega_\rho, \mathcal{R})$ need not be jointly concave in $(\omega_a, \omega_\rho)$, because the cross-derivative is positive—investment in one heterotrophic component enhances the marginal return of the other. This complementarity can generate increasing returns in the heterotrophic subspace, even though each component taken separately is concave.

The interior first-order conditions for the unconstrained objective (ignoring the budget constraint) are

$$L\,\beta\,\omega_p^{\beta-1} = G\,\beta\,\omega_a^{\beta-1}\,\Theta = G\,\omega_a^\beta\,\partial_{\omega_\rho}\Theta = 1,$$

equalizing all marginal returns to unity. If these equations admit a solution with $\omega_p + \omega_a + \omega_\rho < B$, the interior point is a candidate generalist optimum with slack budget. However, because the heterotrophic term has a positive cross-derivative, the full objective is not jointly concave. The Hessian is block-diagonal (since $\omega_p$ is uncoupled), and the determinant of the heterotrophic curvature block is proportional to

$$2\,\kappa\,(1-\beta)\,\omega_\rho\,\mathcal{R} - \beta,$$

so its sign is conditional: it is negative (saddle in the heterotrophic subspace) when $2\,\kappa\,(1-\beta)\,\omega_\rho\,\mathcal{R} < \beta$, and positive when the inequality is reversed. Thus the unconstrained interior point is a saddle whenever sensing investment (or handling time, or resource level) is small enough that the complementarity-induced curvature dominates, but it can be a local maximum when $2\,\kappa\,(1-\beta)\,\omega_\rho\,\mathcal{R} > \beta$. Consequently, no unconditional conclusion follows: depending on parameters, the optimal solution may be an interior generalist with slack budget or lie on the boundary of the budget simplex.

For a binding budget constraint ($\omega_p + \omega_a + \omega_\rho = B$), the appropriate second-order test considers the Hessian restricted to the tangent space $\delta_p + \delta_a + \delta_\rho = 0$. Substituting $\delta_p = -(\delta_a + \delta_\rho)$, the restricted quadratic form is

$$Q = H_p(\delta_a + \delta_\rho)^2 + H_{aa}\,\delta_a^2 + 2\,H_{a\rho}\,\delta_a\,\delta_\rho + H_{\rho\rho}\,\delta_\rho^2,$$

where $H_p < 0$, while $H_{a\rho} > 0$ is the positive cross-derivative. For directions with $\delta_a$ and $\delta_\rho$ of the same sign, the positive cross-term can dominate the negative diagonal terms, making $Q > 0$. Hence the interior stationary point need not be a local maximum even under the binding constraint: when the complementarity dominates, the saddle nature of the heterotrophic subspace persists because the plant direction cannot fully compensate for the positive cross-curvature, and the optimum lies on the boundary; when the diagonal stabilizing curvatures dominate instead, a binding-budget interior generalist can be a local maximum. The global optimum must therefore be determined by comparing fitness values as in the switching logic of (44), numerically in general.

For convex returns ($\beta > 1$), the allocation terms themselves are convex, which generally favors boundary specialization. However, because the Monod response in $\Theta(\omega_\rho, \mathcal{R})$ introduces saturation effects, the full coupled objective need not be jointly convex. Boundary specialization must therefore be verified from the full objective or numerically rather than inferred from the allocation terms alone. Which vertex is optimal follows the same switching logic, with $\mathcal{R}_{\rm sw}$ computed numerically for general $\beta$.

#### Reduction to the two-trait model

If sensing is excluded from the shared budget so that (36) constrains only $\tilde{c}_{\mathcal{L}}\lambda_p + \tilde{c}_{\mathcal{R}}\lambda_a \le B$ while sensing continues to pay its own linear cost $\tilde{c}_\rho \rho$ outside the budget, then for each fixed allocation $\omega_a$ the radius is chosen to maximize the net heterotrophic return

$$G'\,\omega_a^\beta \Theta(\rho, \mathcal{R}) - \tilde{c}_\rho \rho$$

at that resource level, where $G' := \gamma/\tilde{c}_{\mathcal{R}}^\beta$ differs from the shared-budget $G$ of (39) by the factor $\tilde{c}_\rho^{-1}$ absorbed there. Up to the rescaling $L, G', \kappa$ this is exactly the optimization of Section IV, with

$$\Theta(\rho, \mathcal{R}) = \rho\mathcal{R}/(1 + \nu\rho\mathcal{R}), \qquad G' = \gamma/\tilde{c}_{\mathcal{R}}^\beta,$$

and first-order condition

$$\gamma(1-\lambda)^\beta \Theta_\rho = \tilde{c}_\rho \quad (\text{or } G'\omega_a^\beta \Theta_\rho = \tilde{c}_\rho),$$

which gives the same optimal radius $\rho_a$, gain $\Theta_a$ and ecological equilibria.

With $B = 1$ and sensing excluded from the budget, the allocation constraint mirrors, up to the cost weights, the normalized fractions $\lambda + (1 - \lambda) = 1$ of the two-trait model, the only difference being that the shared-budget model caps total spending while the two-trait model charges the same costs linearly without a cap.
